# Neuronal mitochondria transport *Pink1* mRNA via Synaptojanin 2 to support local mitophagy

**DOI:** 10.1101/2021.05.19.444778

**Authors:** Angelika B. Harbauer, Simone Wanderoy, J. Tabitha Hees, Whitney Gibbs, Martha Ordonez, Zerong Cai, Romain Cartoni, Ghazaleh Ashrafi, Chen Wang, Zhigang He, Thomas L. Schwarz

## Abstract

PTEN-induced kinase 1 (PINK1) is a very short-lived protein that is required for the removal of damaged mitochondria through Parkin translocation and mitophagy. Because the short half-life of PINK1 limits its ability to be trafficked into neurites, local translation is required for this mitophagy pathway to be active far from the soma. The *Pink1* transcript is associated with and cotransported with neuronal mitochondria. In concert with translation, the mitochondrial outer membrane protein Synaptojanin 2 binding protein (SYNJ2BP) and Synaptojanin 2 (SYNJ2) are required for tethering *Pink1* mRNA to mitochondria via an RNA-binding domain in SYNJ2. This neuron-specific adaptation for local translation of PINK1 provides distal mitochondria with a continuous supply of PINK1 for activation of mitophagy.

## Introduction

The large size of neurons and the need to maintain mitochondrial health far from the cell body contribute to the vulnerability of neurons to mitochondrial insults (Bolam and Pissadaki, 2012; Harbauer, 2017; Misgeld and Schwarz, 2017). Almost all mitochondrial proteins are encoded in the nucleus. Thus, the replacement and rejuvenation of mitochondria in distal axons and dendrites likely arises from mitochondrial transport in combination with mitochondrial fusion and fission and the mitophagic degradation of damaged organelles. However, there are limitations to the feasibility of transport as the only source of new mitochondrial proteins (Misgeld and Schwarz, 2017). Mitochondria travel at a speed of ∼0.5 µm/s (Misgeld and Schwarz, 2017) and therefore a mitochondrion generated in the cell body will take several days to reach the end of a 1 m long axon. This exceeds the lifetime of many short-lived mitochondrial proteins (Vincow et al., 2013). Likewise, proteins of the respiratory chain encoded by the mitochondrial genome are constantly synthesized in axonal and dendritic mitochondria (Yousefi et al., 2021), but how the complementary nuclear-encoded subunits are supplied is unknown. Local translation of mitochondrial mRNAs provides a possible resolution of this problem. Recent studies have characterized axonal mRNA species (Gumy et al., 2011; Shigeoka et al., 2016; Zivraj et al., 2010), and all such studies have detected a significant amount of nuclear-encoded mitochondrial transcripts in axons. Synaptic synthesis and protein import of mitochondrial proteins has been observed (Cioni et al., 2019; Kuzniewska et al., 2020) and global inhibition of protein synthesis in axons affects mitochondrial health (Hillefors et al., 2007). However, the mechanisms transporting nuclear-encoded mitochondrial mRNAs into neurites are largely unknown.

Failure to induce mitophagy has been linked to the etiology of Parkinson’s disease (PD); two requirements for damage-induced mitophagy – Parkin and PTEN-induced kinase 1 (PINK1) – are mutated in hereditary forms of PD (Exner et al., 2012). This pathway involves the constant, ongoing synthesis, mitochondrial import, and rapid degradation of PINK1 as long as mitochondria are healthy (Narendra et al., 2010). The half-life of PINK1 has been estimated to be in the order of minutes (Ando et al., 2017; Lazarou et al., 2012; Lin and Kang, 2008). Mitochondrial damage stops PINK1 import and the consequent stabilization of PINK1 triggers the translocation of Parkin to this organelle and initiates mitophagy (Durcan and Fon, 2015; Pickrell and Youle, 2015; Yamano et al., 2016). Evidence from our lab determined that acutely damaged mitochondria in axons can undergo local mitophagy and that this mitophagy required both PINK1 and Parkin (Ashrafi et al., 2014). For this mechanism to work in neurons, however, a constant supply of freshly synthesized PINK1 is required regardless of the distance of the mitochondrion from the cell body, thus raising the question of how a nuclear-encoded protein with a half-life of only minutes can be constantly synthesized far from the cell body. Therefore we selected *Pink1* mRNA as a model to investigate the transport of RNA encoding mitochondrial proteins.

Using live-imaging of *Pink1* mRNA, we observed co-transport of *Pink1* mRNA with mitochondria and identified a tethering mechanism that uses the mitochondrial outer membrane protein Synaptojanin 2 binding protein (SYNJ2BP) and a neuron-specific and RNA-binding splice isoform of Synaptojanin 2 (SYNJ2). The neuron-specific co-transport of *Pink1* mRNA with mitochondria enables dendritic and axonal translation of PINK1 to support the removal of damaged organelles and represents a neuron-specific adaptation of the PINK1/Parkin pathway.

## Results

### Local translation is required for PINK1 activity in axons

As the half-life of PINK1 in mammalian neurons may not be the same as that reported in cell lines, we used antibodies to human PINK1 to examine endogenous PINK1 stabilization and clearance in human iPSC-derived neurons (Figure 1A and B). PINK1 levels increased upon depolarization of mitochondria with the ionophore carbonyl cyanide m-chlorophenyl hydrazone (CCCP). This effect could be prevented by addition of the protein synthesis inhibitor cycloheximide (CHX), as expected from the model that calls for continuous synthesis PINK1 in order to respond to mitochondrial damage (Narendra et al., 2010). Upon washout of CCCP, PINK1 levels sharply declined within 30 min to baseline levels, indicating that also in neurons continuous cleavage of PINK1 in polarized mitochondria maintains the protein at low abundance (Fig S1A). Rapid degradation of PINK1 in healthy mitochondria should prevent its effective transport in axons and dendrites. Instead, local translation of *Pink1* mRNA could be the source of the PINK1 protein that supports local mitophagy (Ashrafi et al., 2014). Using a compartmentalized culture system and rodent embryonic hippocampal neurons (Fig. S1B), we asked whether the mitochondrial translocation of Parkin, which is directly dependent on PINK1 stabilization (Ashrafi et al., 2014; Vives-Bauza et al., 2010), occurred in the presence of CHX. CHX was applied to the axonal chamber four hours prior to induction of mitophagy with Antimycin A, using a concentration that has previously been shown to elicit PINK1-dependent mitophagy in axons (Ashrafi et al., 2014). Parkin translocation was visualized by transfection of YFP-tagged Parkin and mitochondrially targeted DsRed (mitoDsRed) (Fig. 1C). The percentage of Parkin-positive mitochondria was quantified before and after mitochondrial depolarization with Antimycin A. Approximately 10% of mitochondria were Parkin positive prior to Antimycin A (Fig. 1C and Ashrafi et al. 2014). Upon addition of Antimycin A, 29% of mitochondria were Parkin-positive. Inhibition of protein synthesis by CHX eliminated 80% of this increase; only 13% of mitochondria accumulated Parkin (Fig. 1D). Thus local translation contributes substantially to efficient induction of mitophagy in axons.

**Figure 1.**
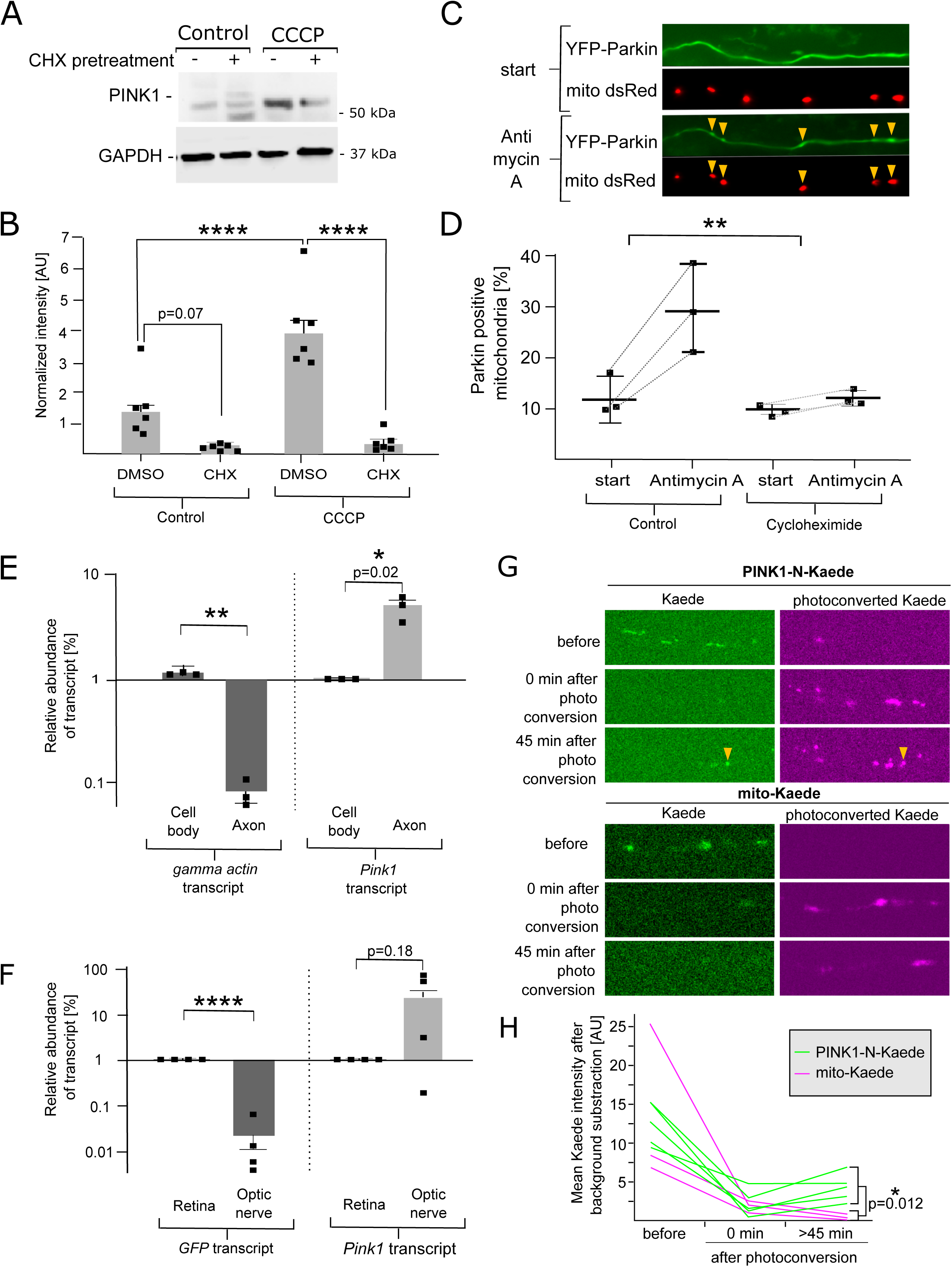
Local translation is required for PINK1 activity in axons. (A) Representative Western blot showing the stabilization of PINK1 in response to the uncoupler CCCP (20 µM 2h) in human iPSC-derived cortical neurons, which can be prevented by addition of 5 µM Cycloheximide (CHX) 20min prior to CCCP. (B) Quantification of PINK1 stabilization as in (A) normalized to GAPDH signal. Data is shown as mean ± SEM; ANOVA with Tukey’s multiple comparisons test, n=6 biological repeats per condition (C) Neurons grown in a compartmentalized culture chamber transfected with YFP-Parkin and a mitochondrial marker were treated with 40 µM Antimycin A. The recruitment of Parkin to mitochondria (yellow arrowheads) was monitored with live cell imaging prior (start) and after 20 min addition of Antimycin A. (D) Quantification of Parkin-positive mitochondria in the axonal compartment before and after Antimycin A treatment in presence or absence of 35 µM Cycloheximide. Data is shown as mean ± SEM; Student’s t-test, n=3 biological repeats per condition. (E) RNA isolated from cell body or axonal compartments of microfluidic devices was analyzed by qPCR. The abundance of the transcripts was normalized to mitochondrial rRNA. Data are shown on a log scale as mean ± SEM; Student’s t-test; n=3 independent microfluidic devices. (F) After intraorbital injection of AAV encoding either GFP or tagged PINK1 transcripts, retinal and optic nerve RNA was collected and the abundance of the exogenous transcripts analyzed by qPCR and normalized to beta Actin. Unlike *GFP* mRNA, *Pink1* mRNA is actively transported into axons. Data are shown on a log scale as mean ± SEM; Student’s t-test, n = 4 retina/optic nerve pairs. (G) Representative images and (H) quantification of the photo-convertible fluorescent protein Kaede fused to PINK1-N (Amino acids 1-225, PINK1-N-Kaede) or Cox8 (Amino acids 1-36, mito-Kaede). The mitochondrial mean signal intensity of the non-photoconverted Kaede was quantified and is shown in the graph. Note that the intensity returning after 45 min post photoconversion is significantly higher in the PINK1-N-Kaede group. Student’s t-test, n = 3-5 axons. p<0.01 (**) and p<0.0001 (****). Scale bars = 10 µm. For further verification of microfluidic devices see Figure S1.

Databases of axonal transcriptomes report *Pink1* mRNA in axons (Shigeoka et al., 2016; Zivraj et al., 2010), which we confirmed by qPCR using hippocampal neurons cultured in microfluidic devices. RNA was extracted from the somal compartment, which represents a mix of neuron cell bodies, dendrites, and axons, as well as some non-neuronal cells, and compared it to RNA extracted from the axon-only compartment. The relative abundances of the *Pink1* transcript and a known cell-body restricted RNA, *gamma actin*, were determined and normalized to the amount of mitochondrial rRNA, which is transported inside mitochondria as part of mitochondrial trafficking and therefore not excluded from the axon. The *gamma actin* transcript was almost absent from the axonal fraction, whereas *Pink1* mRNA was readily detected (Fig. 1E and S1C).

To establish whether *Pink1* mRNA is also present in adult axons *in vivo*, we used adeno-associated virus injection into the eye to express transcripts for either a sequence-tagged *Pink1* or *GFP* control transcript in mouse retinal ganglion cells, cells which project their axons to the brain exclusively through the optic nerve. Four weeks after the injection, RNA preparations from retina and from the optic nerve segment prior to the optic chiasm were analyzed by qPCR and the observed amounts normalized to the abundance of *beta actin* transcript to account for variances in the amount of tissue harvested. The exogenous *Pink1* transcript was detectable in both tissues and had therefore been transported into the optic nerve, whereas the control *GFP* transcript was barely detectable in the axon (Fig. 1F). The selective presence of the expressed *Pink1* mRNA relative to the *GFP* mRNA argues against simple diffusion as a mechanism of transport and corroborates our findings of axonal *Pink1* mRNA in an adult *in vivo* model.

In order to test whether we could observe translation of PINK1 in axons, we used the photoconvertible protein Kaede and targeted it to mitochondria using either a generic mito-targeting domain (mito-Kaede, using the COX8a presequence, amino acids 1-36) or a portion of the PINK1 protein (PINK1-N-Kaede, amino acids 1-225). By repeated illumination with 405 nm laser light, Kaede is irreversibly converted into a red fluorophore, which allows the detection of newly synthesized, non-photoconverted and therefore green-fluorescent Kaede (Raab-Graham et al., 2011). The N-terminus of PINK1 was sufficient to support local translation, as we were able to detect freshly synthesized, non-photoconverted signal for the PINK1-N-Kaede fusion protein 45 min after photoconversion in axonal mitochondria that were also positive for the photoconverted Kaede-red, whereas the signal for mito-Kaede did not recover (Fig. 1G and H). This correlates with the sequences within the transcript that are necessary for transport of the *Pink1* transcript as will be shown subsequently. The lack of recovery for mito-Kaede argues that a transcript not explicitly targeted to neurites could not supply the distal axons in this short time frame via protein transport from the soma, further strengthening the hypothesis that local translation supplies a substantial fraction of fresh PINK1 protein.

### *Pink1* mRNA localizes to mitochondria in neurons

The endogenous *Pink1* transcript, as localized by RNAscope *in situ* hybridization, occurs in clusters along axons that are reminiscent of the distribution of mitochondria (Fig. 2A). Indeed, immunolabelling of the mitochondrial protein ATP5a in the same axons, revealed extensive co-localization of the two signals (Fig. 2A, left panel). *Pink1* transcripts also localized to dendritic mitochondria (Fig. 2A right panel). With STED imaging, the *Pink1* mRNA signal was resolved into multiple small puncta on each mitochondrion. In contrast, the control *beta actin* transcript did not overlap with axonal mitochondria (Fig. 2B).

**Figure 2.**
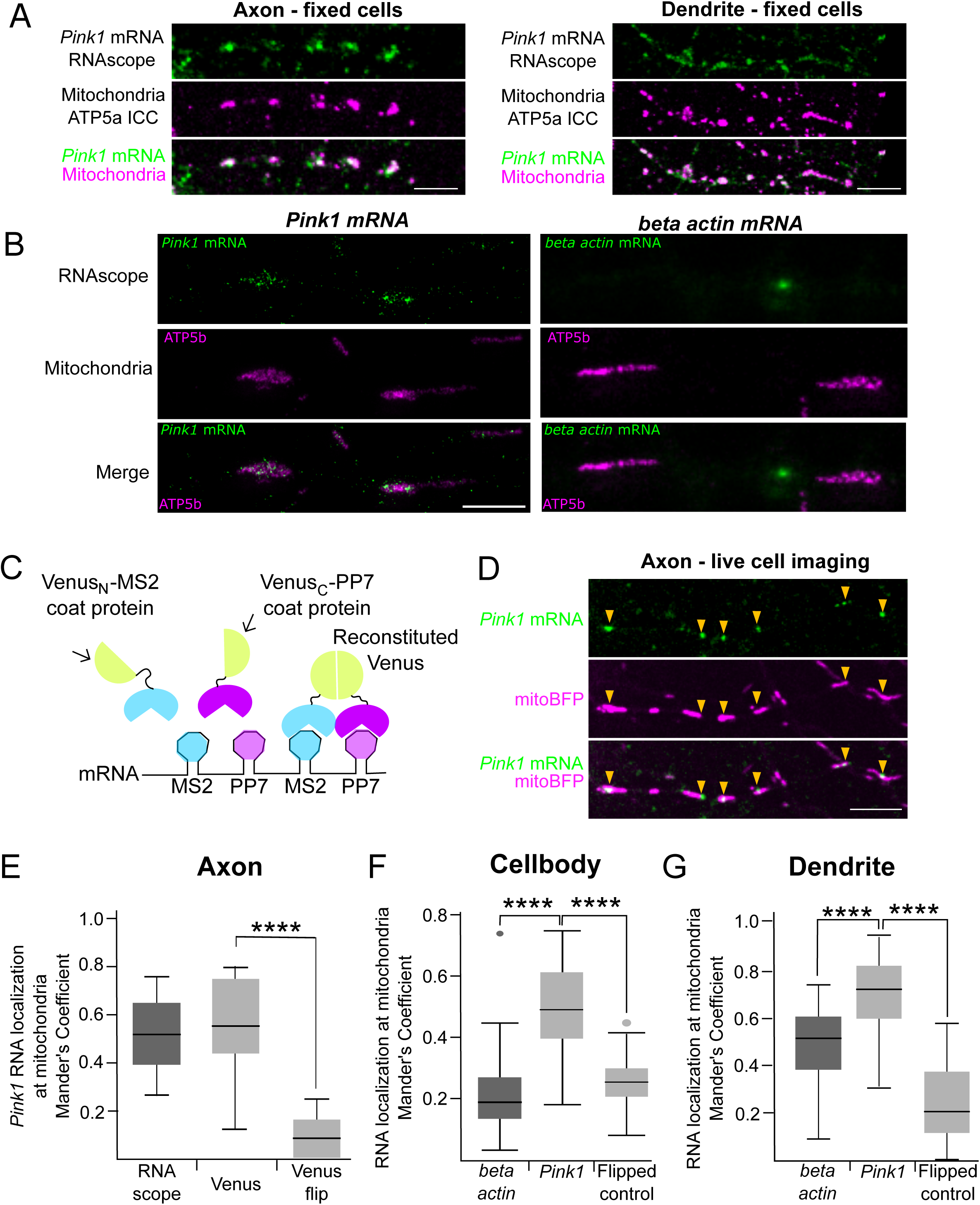
*Pink1* mRNA localizes to mitochondria in neurons. (A) RNAscope *in situ* hybridization reveals *Pink1* mRNA localization to mitochondria in axons and dendrites. Scale bar = 10 µm. (B) Representative superresolution STED images for endogenous *Pink1* and *beta actin* RNA by *in situ* hybridization (RNAscope) and mitochondria detected by immunostaining for ATP5b. Scale bar = 2 µm. (C) Schematic of MS2/PP7-split-Venus method for mRNA imaging. (D-G) Live cell imaging of *Pink1* and *beta actin* RNA in hippocampal neurons using the MS2/PP7-split-Venus method. (D) Representative image of co-localization of the tagged RNA with mitochondrially targeted BFP (mitoBFP). Scale bar = 10 µm. (E) Mander’s correlation coefficient for RNA and mitochondrial channels. “Venus-flip” indicates that the mitochondrial channel, after digital straightening of the axon, had been flipped horizontally before quantification. Student’s t-test, n = 10 axons; p<0.0001 (****). (F and G) Mander’s correlation coefficient analysis between RNA and mitochondrial channels in cell bodies and dendrites. “flipped control” indicates that the mitochondrial channel, after digital straightening in the case of dendrites, had been flipped horizontally before quantification Student’s t-test, n = 39-43 cell bodies, 16-20 dendrites; p<0.0001 (****). Scale bar = 10 µm. For representative images, colocalization with endosomes and further detail on the MS2/PP7-split-Venus method see Figure S2.

To visualize transport of *Pink1* mRNA in live neurons, we used the MS2/PP7-split-Venus approach for background-free mRNA imaging (Wu et al. 2014, Fig. 2C). This method combines the specific binding of the MS2 or PP7 viral coat proteins to their respective RNA stem loops with fluorophore reconstitution. Briefly, the mRNA of interest is expressed carrying a tag consisting of alternating MS2 and PP7 stem loops. In the same cell, the PP7- and MS2-binding coat proteins are expressed and each is fused to one half of split-Venus. Therefore, only in the presence of the transcript are the two halves of Venus in sufficient proximity to reconstitute a fluorescent protein (Fig. 2C). We therefore added 12 copies of PP7 and MS2 stem loops to a rat *Pink1* construct that included the relatively short 3’ and 5’ untranslated regions (UTR) of this gene. As the split-Venus approach contains the entire open reading frame (ORF), expression of this construct will lead to overexpression of the PINK1 protein. In order to avoid confounding effects of PINK1 overexpression, which for example, induces mitochondrial arrest (Wang et al., 2011), we also introduced a kinase dead mutation (K219M). The stem/loop-tagged *Pink1* mRNA was readily detected in cell bodies and dendrites (Fig. S2A). However, detection of the transcript in axons (marked by AnkyrinG-mCherry) was limited using this method due to the difficulty of having adequate axonal levels of the fluorescently-tagged coat proteins present in axons (Fig. S2B and C). Nevertheless, small *Pink1* mRNA puncta were observed in some proximal axons (Fig. 2D). Unlike the RNAscope data, in which every mitochondrion was associated with detectable *Pink1* mRNA, the reconstituted Venus fluorescence was not detected on every mitochondrion, which likely represents the technical limit to the sensitivity of the method as applied to neurites.

Whether detected by RNAscope or with the MS2/PP7 split-Venus method, the *Pink1* transcripts were on mitochondria (Fig. 2A, B and D). Using the Mander’s correlation coefficient we quantified our imaging results and found a significant overlap between the *Pink1* mRNA and mitochondrial signals regardless of the method used (Fig. 2E). In order to exclude that the relative frequency of mitochondria in axons would produce the same overlap randomly, we flipped the mitochondrial channel of the straightened image and performed the same quantification. The mitochondrial localization of the *Pink1* mRNA was significantly higher in the original images compared to the flipped control (Venus-flip, Fig. 2E).

As seen with RNAscope, *Pink1* mRNA visualized with MS2/PP7 split-Venus also co-localized with dendritic mitochondria and to a somewhat lesser extent with mitochondria in the cell body. We observed a significantly higher overlap in all cases when comparing the Mander’s correlation coefficients in neuronal compartments as compared to the flipped quantification (Fig 2F-G and S2D).

Additionally, when *beta actin* mRNA was similarly tagged, small fluorescent puncta were observed, as expected for a transcript transported as a ribonucleoprotein particle (RNP). The *beta actin* transcripts did not co-localize with mitochondria in the cell body (Mander’s coefficient was not significantly higher than the flipped quantification control), but we observed a marginal increase in colocalization in dendritic mitochondria. The slightly higher Mander’s coefficient in dendrites is in line with observations that also non-mitochondrial transcripts including *actin* may use mitochondria as platforms for protein synthesis (Cioni et al., 2019; Spillane et al., 2013). Likewise, the *beta actin* mRNA has been found to localize to mitochondria in HeLa cells (Briley et al., 2015), supporting the physiological relevance of a closer association between mitochondria and this transcript.

Nonetheless, a significant difference remained between the two transcripts indicating a high likelihood that, unlike *beta actin* mRNA, *Pink1* transcripts are actively localized to mitochondria (Fig. 2E-G). Additionally, we tested other mitochondrial transcripts. We chose two transcripts of MTS-containing proteins that have previously been reported to be either axonally localized, *Cox4i* (Kar et al., 2017), or mitochondrially localized, *Atp5f1b* (Margeot et al., 2002). Both transcripts are also detected in axonal transcript databases (Gumy et al., 2011; Shigeoka et al., 2016), yet encode proteins of very different half-lives (COX4i: approx. 30 min, ATP5F1b: approx. six days) (Schwanhäusser et al., 2011). Both the full length *Cox4i* and the *Atp5f1b* transcripts (including their 3’ and 5’UTRs) were significantly colocalized with mitochondria relative to the respective flipped-image control, albeit to a lesser extent compared to *Pink1* (Fig. S4E and F). This is in line with the findings by Cohen et al. in the accompanying manuscript (Cohen et al., 2021), that also find extensive co-localization of mitochondrial transcripts with neuronal mitochondria.

As late endosomes had recently been implicated in the translation of mitochondrial transcripts in axons (Cioni et al., 2019), we also analyzed *Pink1* mRNA localization in relation to endosomes labelled by mCherry-Rab7a. In the cell body the *Pink1* transcript did not significantly overlap with late endosomes (Fig. S2G and H), whereas its association with mitochondria in the soma was clear. This colocalization in the soma together with the presence of the transcript on mitochondria in processes suggests that the *Pink1* transcript is transported through its association with mitochondria rather than in separate RNPs.

### *Pink1* mRNA is co-transported with mitochondria

Because *Pink1* mRNA was present on mitochondria in both axons and dendrites but live imaging of *Pink1* particles in axons was limited, we analyzed the transport of the mRNA mainly in dendrites. In time-lapse images and kymographs, *Pink1* mRNA was on both moving and stationary mitochondria (Fig. 3A; movie S1). Most *Pink1* mRNA particles and their associated mitochondria were stationary, as expected from the predominance of stationary mitochondria in dendrites (Overly et al., 1996) (red arrowhead, Fig. 3A). *Pink1* mRNA particles were also present on moving mitochondria (yellow arrowhead, Fig. 3A) and their movements mirrored those of the organelles, as indicated by their overlapping kymography traces (Fig 3A, lower panel). The *Pink1* mRNA and mitochondria behaved as if bound together; the *Pink1* mRNA stopped and started when its associated mitochondrion stopped and started. On a few occasions, we observed *Pink1* mRNA particles that were not on a detectable mitochondrion. In some of these cases, kymographs suggested possible associations and dissociations from moving mitochondria (Fig 3B, S3A; movie S2). In most cases, however, the mRNA and mitochondrion remained together for the duration of the observation and this was true in both dendrites and axons (Fig. S3B; move S3). In 43 movies from dendrites we observed 96 moving mRNA particles, and in 89% of the cases *Pink1* mRNA and mitochondria moved in synchrony (compare Fig. 3A-C; movies S1-3). In cases when we could not detect a corresponding mitochondrial trace the mRNA particles were stationary or moved only very short distances, while all long-range transport events happened in association with mitochondrial transport (Fig. 3B, C, S3; movie S2). There was no obvious directional preference of movement (time spent in motion for *Pink1* mRNA particles: anterograde moving 3.13% ±0.53%; retrograde moving 2.51% ±0.51%; p=0.41, Student’s t-test, n=29 dendrites). This implies that mitochondrial transport of *Pink1* mRNA is not directional and strictly for the delivery of this transcript from the soma to distal regions. Rather the mRNA movement mirrors the bidirectional movement of mitochondria (Overly et al., 1996). Together with the presence of the mRNA on the stationary pool of mitochondria, the bidirectional movement implies an ongoing association even in the periphery of the cells.

**Figure 3.**
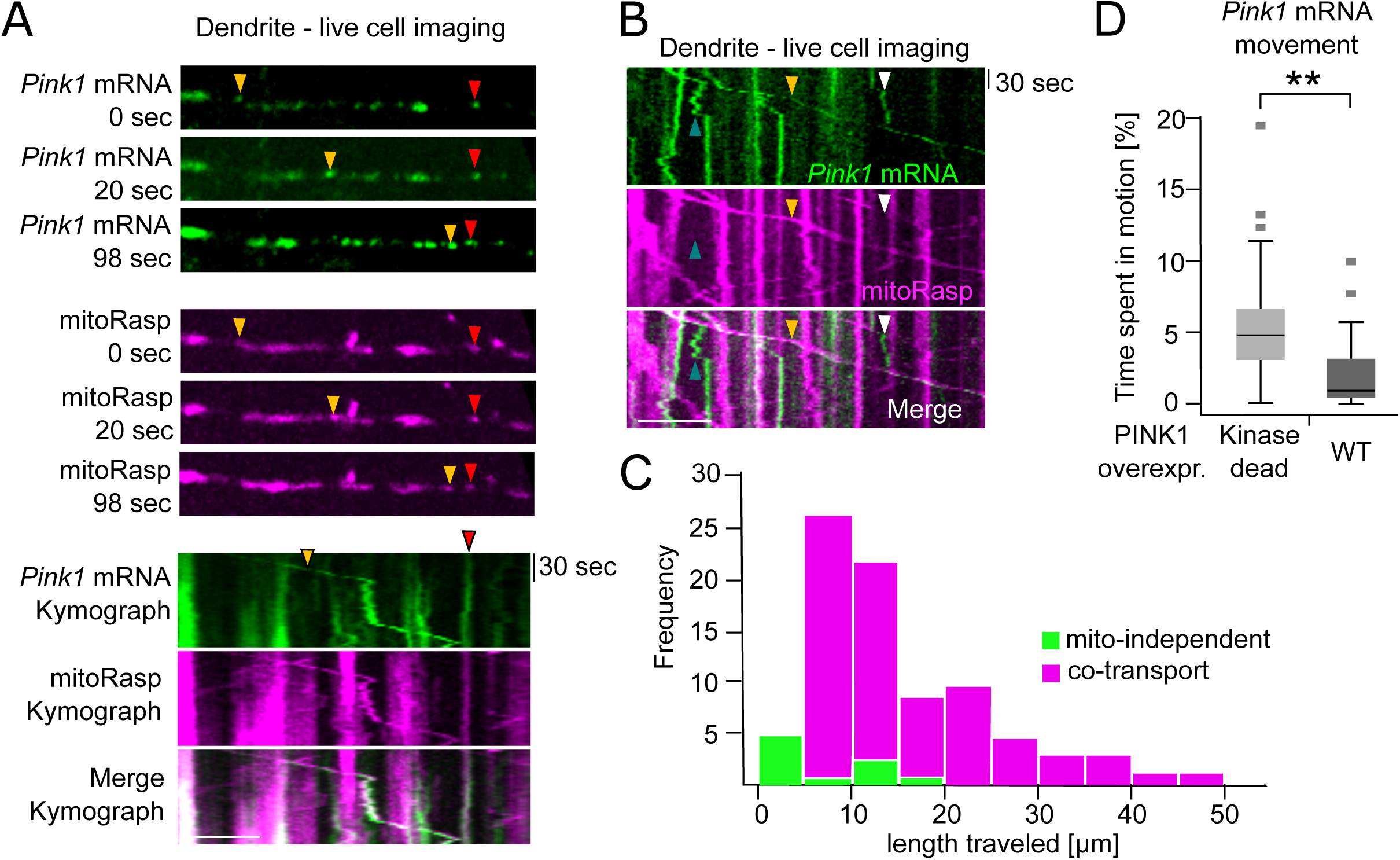
*Pink1* mRNA is co-transported with mitochondria. (A) Still-images and kymograph of a time-lapse movie from a dendrite showing *Pink1* mRNA co-transported with a mitochondrion (yellow arrowhead) as well as stationary mitochondria associated with a *Pink1* mRNA particle (red arrowhead). Scale bar = 5 µm (B) Kymograph in which *Pink1* mRNA appeared to transiently occur without an associated mitochondrion. While most mRNA particles were associated with mitochondria, including during transport (yellow arrowhead), particles without a mitochondrion were occasionally seen and might undergo short-range independent movement (blue and white arrowheads). A potential interpretation of this kymograph as indicating association and dissociation of the *Pink1* mRNA from moving mitochondria is schematized in Figure S3. Scale bar = 10 µm. (C) Histogram depicting frequency of observed movements of 96 moving *Pink1* mRNA particles from 46 dendrites over the indicated distances. (D) Overexpression of wild-type (WT) PINK1 protein decreases *Pink1* mRNA motility relative to kinase-dead PINK1 protein. Average time spent in motion per dendrite was analyzed in n = 26-29 dendrites from three independent experiments. Student’s t-test, p<0.01 (**).

Overexpression of catalytically active PINK1 protein inhibits mitochondrial movement (Wang et al., 2011). We therefore compared the motility of the mRNA encoding wild-type (WT) and kinase-dead PINK1, to determine the consequences of the resulting overexpressed proteins. As expected from the extensive coupling of *Pink1* mRNA transport to mitochondria, the transport of the mRNA encoding the catalytically active version was affected by the resulting overexpression of catalytically active PINK1 (Fig. 3D). On average, *Pink1* mRNA particles spent 60% less time in motion, when the WT form of PINK1 protein was expressed compared to the expression of the kinase-dead mutant.

### Translation of the PINK1 mitochondrial targeting sequence is necessary but insufficient for *Pink1* mRNA association with mitochondria

To identify the mechanism that tethers the *Pink1* transcript to the mitochondrial surface, we expressed shortened versions of the *Pink1* transcript fused to the coding sequence for *BFP* (Fig. 4). The *BFP* sequence alone did not associate with mitochondria. We therefore asked whether any portions of the *Pink1* transcript could convey mitochondrial association onto a chimeric construct (Fig. 4A). We analyzed the extent of mitochondrial transcript colocalization in neuronal somata with the MS2/PP7-split-Venus system. Although many RNA-binding proteins (RBPs) bind within the untranslated regions (UTRs) of mRNAs (Holt and Schuman, 2013), neither of the *Pink1* UTRs was sufficient for inducing the mitochondrial localization of the *BFP* transcript (Fig. 4A and S4A). Likewise, a transcript that included the C-terminal part of the PINK1 protein (nucleotides 676-1743 of the open reading frame (ORF) and the 3’ UTR of *Pink1*) did not convey localization to mitochondria (Fig. 4A and S4A). However, inclusion of the N-terminal part of the PINK1 protein (the 5’ UTR and nucleotides 1-675 of the *Pink1* ORF) was sufficient to localize the mRNA to mitochondria (*Pink1 5’UTR+N-BFP*, Fig. 4A and B). This is consistent with our observation that fusion of the PINK1 N-terminus (amino acids 1-225, encoded by bp 1-675) supports local translation as measured by Kaede reappearance after photo-conversion (Fig. 1G and H).

**Figure 4.**
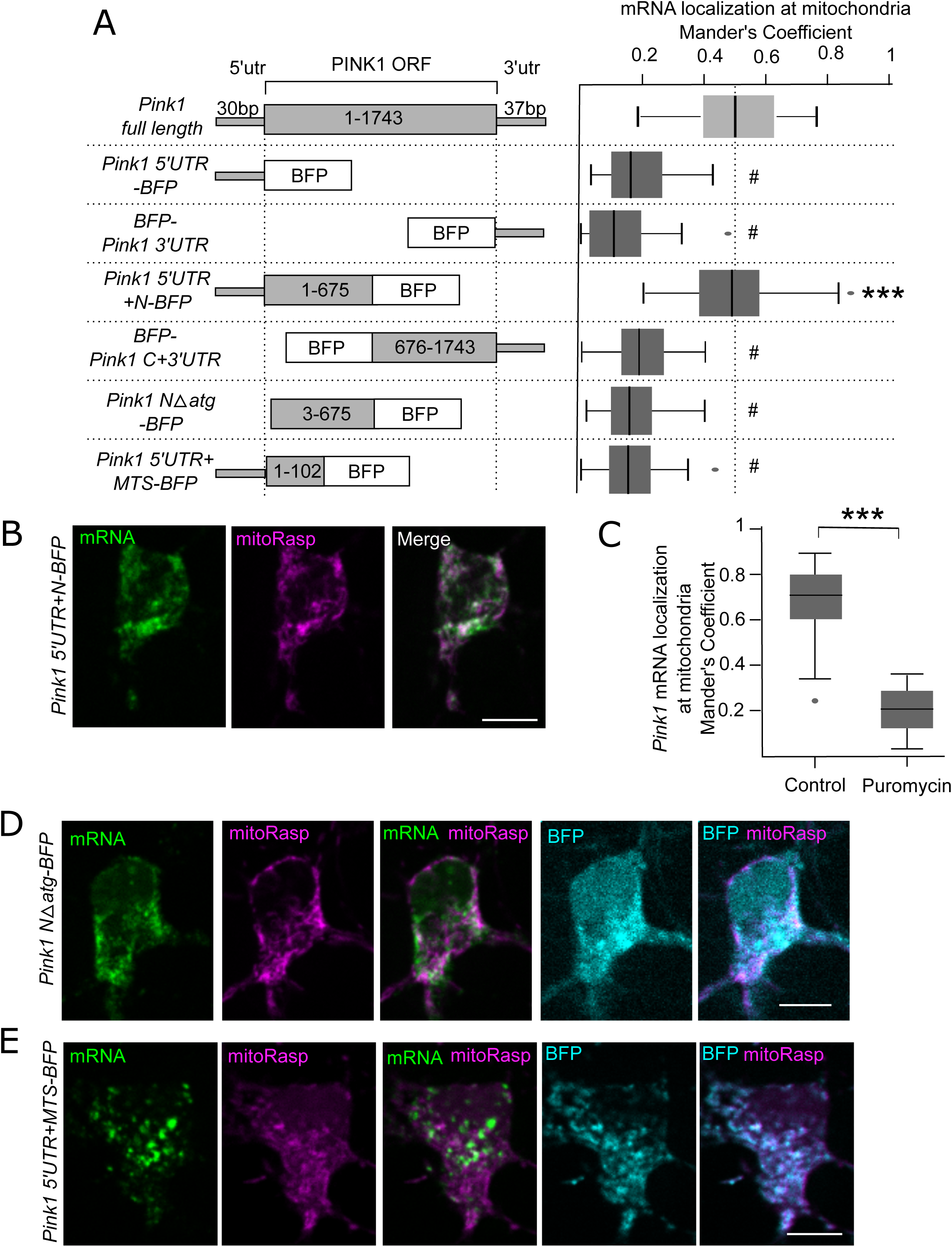
Translation of the PINK1 mitochondrial targeting sequence is necessary but insufficient for *Pink1* mRNA association with mitochondria. (A) Schematic representation of constructs used and Mander’s correlation coefficients between the indicated *Pink1/BFP* chimeric constructs and mitochondria in hippocampal cell bodies. The distribution of full length *Pink1* was repeated from Figure 2F for comparison. One-way ANOVA with Bonferroni post-hoc test; n = 28-41 cell bodies, p<0.001 (***), p>0.05 (#). (B) Representative images for *Pink1 5’UTR+N-BFP*. (C) Mander’s correlation coefficient between *Pink1* transcript (kinase dead) and mitochondria in the presence or absence of puromycin. Student’s t-test, n = 26-28 cell bodies, p<0.001 (***). (D) Representative images of *Pink1-N-Δatg-BFP* RNA and protein. (E) Representative images for *Pink1 5’UTR+MTS-BFP.* Please note that the *Pink1 5’UTR+MTS-BFP* mRNA is largely cytosolic, although the encoded BFP protein localizes exclusively to mitochondria. Scale bars = 10 µm. For representative images see Figure S4.

The *Pink1 5’UTR+N-BFP* construct contained both the start codon and the mitochondrial targeting sequence (MTS) of PINK1 and thereby suggested that translation of the transcript might be required for its mitochondrial localization. Indeed, expression of the full length *Pink1* construct in the presence of the translation inhibitor puromycin altered the *Pink1* mRNA distribution. After one hour incubation with 200 µg/ml puromycin, full length *Pink1* mRNA had shifted from mitochondria to the cytosol (Fig. S4B, quantified in Fig. 4C). Puromycin dissociates both nascent chains and mRNA from ribosomes, thus translation is required for targeting the *Pink1* mRNA to mitochondria. A construct lacking the start codon did not localize to mitochondria (*Pink1 NΔatg-BFP*, Fig. 4A and D), which further corroborated the requirement for translation. We still observed translation of the BFP portion of this transcript (Fig. 4D), presumably due to a downstream methionine.

The requirement for translation raised the possibility that the association of the mRNA with mitochondria was driven by the MTS on the nascent chain, similar to the targeting of ribosomes to the ER upon translation of ER proteins (Hegde and Bernstein, 2006). Although mitochondria can import proteins after their translation, co-translational import may occur as soon as the MTS leaves the ribosomal tunnel (Verner, 1993). The nascent MTS interacts with the receptors of the translocase of the outer membrane (TOM) complex (Harbauer et al., 2014), thereby driving the association of the ribosome and mRNA with the mitochondrial surface. This phenomenon has been observed in yeast and was recently also demonstrated in cultured mammalian cells (Fazal et al., 2019; Vardi-Oknin and Arava, 2019). We therefore hypothesized that the MTS of PINK1 might be sufficient to localize its mRNA to mitochondria. The first 34 amino acids of PINK1 are sufficient to drive protein import into mitochondria (Okatsu et al., 2015), and indeed when we expressed a construct consisting of the 5’UTR and MTS of *Pink1* fused to *BFP* (*Pink1 5’UTR+MTS-BFP*), the BFP-fusion protein localized to mitochondria (Fig. 4E). This chimeric *Pink1/BFP* transcript, however, was still cytoplasmic (Fig. 4E). The interaction of the nascent chain with the TOM complex therefore is not sufficient to stabilize the transcript on mitochondria. The Mander’s correlation coefficient was comparable for the *Pink1 5’UTR+MTS-BFP* and *Pink1 NΔatg-BFP* constructs (Fig. 4A). As these two overlapping transcripts together include all the sequences present in the mitochondrially targeted *Pink1 5’UTR+N-*BFP construct, yet neither is sufficient to promote mitochondrial localization on its own, multiple sequences within the mRNA seem required for efficient mitochondrial localization. This prompted us to investigate the involvement of RBPs in *Pink1* mRNA localization.

### SYNJ2BP knock down redistributes *Pink1* mRNA into RNA granules and inhibits local mitophagy

Overexpression of the mitochondrial outer membrane protein SYNJ2 Binding Protein (SYNJ2BP, also known as Outer Membrane Protein 25, OMP25) leads to an increase in mito-ER contact sites that are enriched for ribosomes (Hung et al., 2017). A recent preprint now also connects SYNJ2BP with the localization of mitochondrial RNAs to the outer mitochondrial membrane (Qin et al., 2020).

We therefore knocked down SYNJ2BP expression in hippocampal neurons (Fig. S5A) and analyzed the localization of the *Pink1* transcript using the MS2/PP7-split-Venus method. Knockdown of SYNJ2BP, but not control shRNA, diminished the interaction between *Pink1* mRNA and mitochondria in the cell body and prevented the transport of the transcript into dendrites (Fig. 5A and B). This effect was specific to the loss of SYNJ2BP as expression of an shRNA-resistant SYNJ2BP rescued this effect (Fig 5B). The *Pink1* mRNA left in the cell body formed larger aggregates that colocalized with RFP-DDX, an RNA helicase that is present in processing bodies (P-bodies, Fig. 5C and S5B). P-bodies are a cellular RNA storage compartments that sequester mostly untranslated mRNA and contain many RNA degrading enzymes (Sfakianos et al., 2016; Wilbertz et al., 2019). In contrast to its effect on *Pink1* mRNA, the colocalization of *beta actin* mRNA and RFP-DDX6 was unchanged by SYNJ2BP knock down (Fig. S5C and D). These results indicate that loss of SYNJ2BP has a selective effect on *Pink1* mRNA localization.

**Figure 5.**
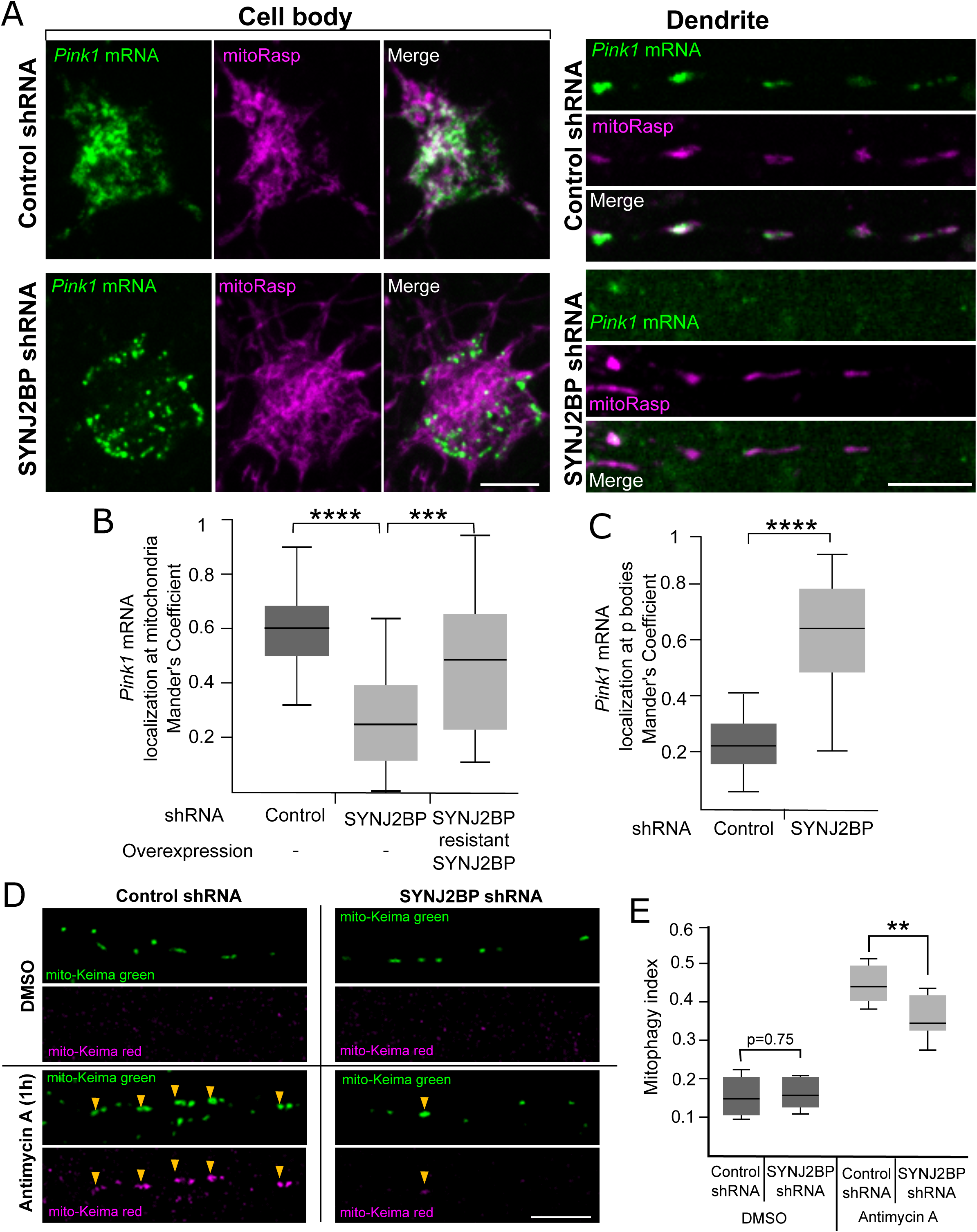
SYNJ2BP knock down redistributes *Pink1* mRNA to RNA granules and inhibits local mitophagy. (A) Neurons were treated with either control or SYNJ2BP shRNA for imaging of *Pink1* transcripts (kinase dead) by the split-Venus method and mitochondria. Representative images from the cell bodies and dendrites are shown. (B) Colocalization in the cell body as quantified with Mander’s correlation coefficient. Coexpression of an shRNA-resistant SYNJ2BP restored localization of *Pink1* transcripts to mitochondria. Student’s t-test, n = 23-31 cell bodies, p<0.0001 (****). (C) Mander’s correlation coefficient for colocalization in cell bodies of *Pink1* transcript and P-bodies marked with RFP-DDX. Student’s t-test, n = 16-20 cell bodies, p<0.0001 (****). (D) Representative images of the pH-sensitive fluorophore mKeima targeted to mitochondria in axons from neurons also expressing either Control or SYNJ2BP shRNA, with and without Antimycin A treatment. (E) Mitophagy index for neurons treated as in (D). Student’s t-test, n = 9-10 axonal field of views, p<0.01 (**). Scale bars = 10 µm. For the localization of *beta actin* mRNA and validation of the shRNA please refer to Figure S5.

As RNA localization to P-bodies may be linked to its degradation we analyzed the levels of *Pink1* mRNA upon SYNJ2BP knockdown but did not find a decrease, but rather a trend towards an increase in the amount of *Pink1* transcript (Fig. S5E), which may be due to compensatory mechanisms upregulating *Pink1* expression. Also, SYNJ2BP knockdown did not prevent expression of an mRNA that would otherwise be targeted to mitochondria; we still observed translation of the PINK1-BFP chimeric construct (Fig. S5F) in the cell body.

The requirement for SYNJ2BP for *Pink1* mRNA association with mitochondria and cotransport predicted that knockdown of SYNJ2BP would prevent local activation of axonal mitophagy, in keeping with the dependence of the PINK1/Parkin pathway on local axonal translation (Fig. 1). To measure mitophagy when SYNJ2BP is knocked down, we turned to the mitochondrially localized, pH-sensitive reporter mito-mKeima (Katayama et al., 2011), a fluorophore with a bimodal excitation spectrum depending on protonation of its chromophore. At neutral pH, when mitochondria are healthy, Keima is most effectively excited at 442 nm (represented in green), whereas under acidic conditions, upon induction of mitophagy and fusion with lysosomes, Keima excitation peaks at 550 nm (pseudocolored red). Consistent with the need for *Pink1* mRNA transport, knockdown of SYNJ2BP reduced the incidence of Antimycin A-induced mitophagy in distal axons (Fig. 5D and E), confirming the local loss of PINK1 function.

### *Pink1* mRNA mitochondrial localization is neuron-specific and depends on SYNJ2a

*Pink1* mRNA was not on mitochondria in COS-7 cells (Fig. 6A); the mitochondrial association of the transcript may therefore be neuron-specific. The lack of mitochondrial localization of the transcript allowed us to ask what missing neuronal component was needed to bring the *Pink1* mRNA to COS-7 mitochondria. As SYNJ2BP is ubiquitously expressed, we analyzed if its interaction partners, the SYNJ2a splice variant of Synaptojanin 2 (SYNJ2) (Nemoto and De Camilli, 1999) or the ER protein Ribosome Binding Protein 1 (RRBP1)(Hung et al., 2017) are enriched in neurons over fibroblasts. *Synj2a* transcripts were five-fold higher in hippocampal cultures than in fibroblasts (Fig. 6B), unlike the *Rrbp1* transcripts levels, which were seven-fold higher in fibroblasts than neurons (Fig. 6C).

**Figure 6.**
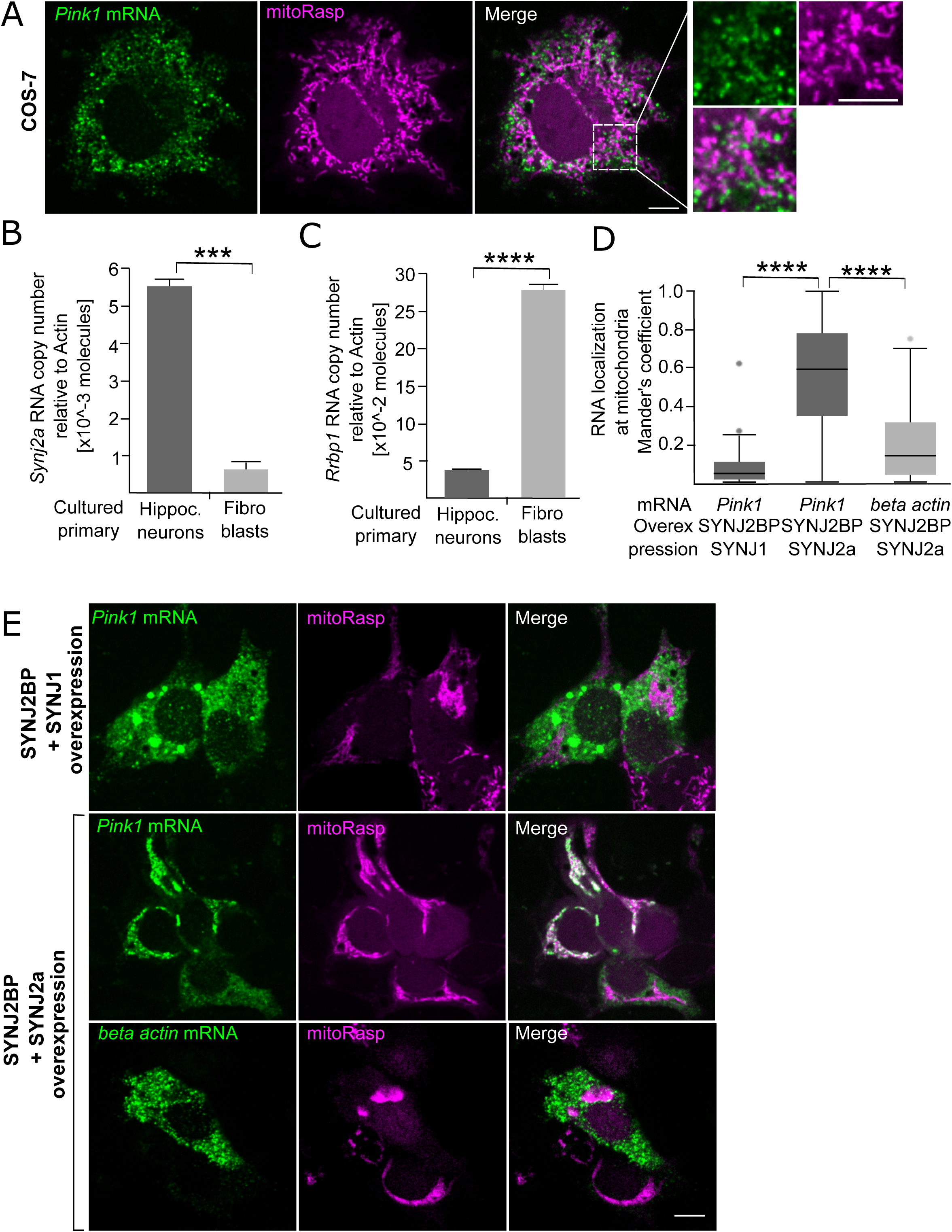
*Pink1* mRNA localization to mitochondria is neuron specific and depends on SYNJ2a. (A) *Pink1* transcript is not localized to mitochondria in COS-7 cells. (B) qRT-PCR from primary fibroblasts and hippocampal neurons comparing the expression levels of the *Synj2a* splice variant. Data are shown as mean ± SEM; Student’s t-test; n=3-5 cultures. (C) qRT-PCR from primary fibroblasts and hippocampal neurons comparing the expression levels of the *Rrbp1* transcript. Data are shown as mean ± SEM; Student’s t-test; n=3 cultures. (D) Mander’s Coefficients of *Pink1* and *beta actin* RNA colocalization with mitochondria in COS-7 cells overexpressing the indicated proteins. Data are shown as mean ± SEM; Student’s t-test; n ≥ 3 experiments scoring ≥30 cells per condition total. p<0.01 (**). (E) Representative images of *Pink1* and *beta actin* transcript localization in COS-7 cells with overexpression of the indicated proteins. Scale bars = 10µm. For knock down of SYNJ2 in neurons, please refer to Figure S6.

We therefore overexpressed SYNJ2a and SYNJ2BP in COS-7 cells and found that *Pink1* mRNA was thereby relocalized to mitochondria (Fig. 6D and E). In contrast, overexpression of the related protein Synaptojanin 1 (SYNJ1) together with SYNJ2BP did not change the cytoplasmic localization of the *Pink1* transcript. The effect of SYNJ2a overexpression was specific for *Pink1* mRNA; *beta actin* mRNA remained cytosolic (Fig. 6D and E). As had been reported in Nemoto and De Camilli (1999) overexpression of SYNJ2BP and SYNJ2a also led to mitochondrial clustering (compare Fig. 6E).

Given our findings in COS-7 cells, we tested the importance of SYNJ2 for the mitochondrial localization of *Pink1* mRNA in neurons. Localization of *Pink1* mRNA with neuronal mitochondria was diminished by knock down of SYNJ2, but rescued by co-expression of an shRNA-resistant SYNJ2 construct (Fig. S6A and B). We therefore conclude that in addition to co-translational targeting, SYNJ2 acts as a mitochondrial anchor for *Pink1* mRNA.

### RNA-binding by SYNJ2 is necessary for *Pink1* mRNA localization to mitochondria

SYNJ2 is an inositol 5’-phosphatase with an established role in endocytosis (Nemoto et al., 1997). It is, however, also predicted to contain an RNA recognition motif (RRM) domain (Hsu and Mao, 2015), although its ability to bind nucleotides has not been established. Based on the homology between different RRM domains (Maris et al., 2005), we identified the residues Valine 909, Glutamine 951 and Leucine 953 (VQL) in SYNJ2 as likely to be involved in RNA binding. We expressed the myc-tagged RRM domain of rat SYNJ2 in HEK cells and used 254 nm UV-Crosslinking and Immuno-Precipitation (CLIP) to determine if the RRM domain could be crosslinked to RNA (Castello et al., 2012; Greenberg, 1979). Upon UV irradiation, we observed several crosslinked species of WT SYNJ2 RRM running at higher molecular weights, as expected for an RNA-binding domain (Fig. 7A). Mutation of the VQL residues to Alanine abolished its RNA-binding ability and prevented the appearance of the cross-linked species (VQL/AAA, Fig. 7A lane 3).

**Figure 7.**
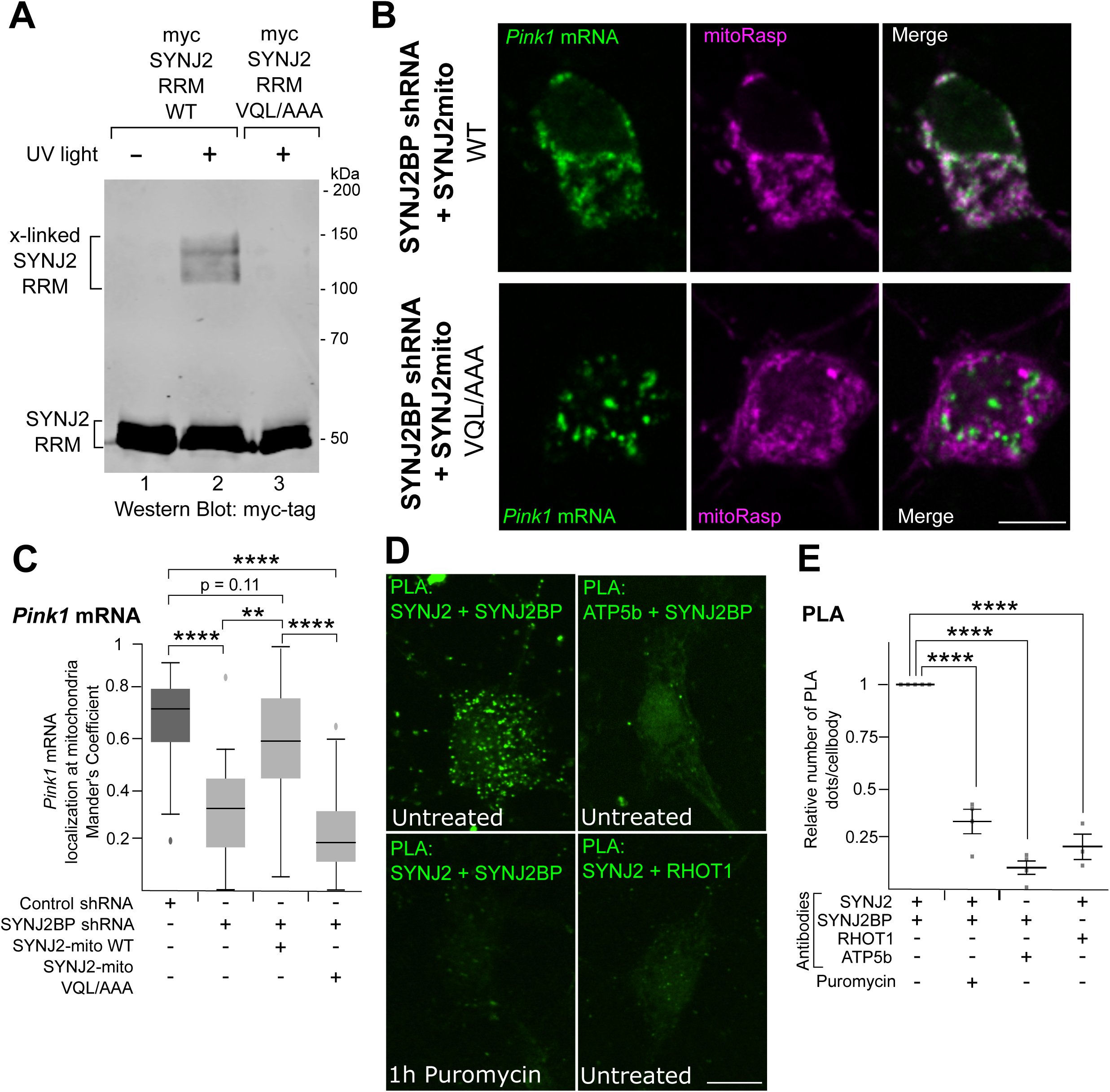
RNA-binding by SYNJ2a is necessary for *Pink1* mRNA localization to mitochondria. (A) Myc-tagged SYNJ2-RRM constructs were expressed in HEK 293T cells and irradiated as indicated with 254 nm UV light. Lysates were immunoprecipitated with anti-myc and a representative anti-myc Western blot is shown. (B) Representative images of neuron cell bodies expressing SYNJ2BP shRNA and either WT or VQL/AAA SYNJ2mito, in addition to the split-Venus reporter for *Pink1* mRNA and mitoRasp. Scale bar = 10 µm. (C) Colocalization as quantified with Mander’s correlation coefficient between mitochondria and *Pink1* (kinase dead) transcripts for cells as in (B). Student’s t-test, n = 20-23 cell bodies; p<0.01 (**), p<0.0001 (****). (D) Representative images displaying the proximity ligation assay (PLA) using the indicated antibodies in the presence or absence of puromycin. (E) Quantification of PLA results as in (D) Student’s t-test; n ≥ 3-5 experiments scoring ≥15 cells per condition total. p<0.0001 (****). Data are shown as mean ± SEM; scale bars = 10 µm. For *beta actin* imaging in the presence of either WT or VQL/AAA SYNJ2mito in SYNJ2BP shRNA treated neurons see Figure S7.

To determine if the RNA-binding property of SYNJ2a was sufficient to target *Pink1* mRNA to mitochondria, we constructed both a WT and a VQL/AAA version of an artificial tether analogous to the constructs used in Nemoto *et al*., 1997. By directly fusing the open reading frame of SYNJ2a to the transmembrane domain of SYNJ2BP (SYNJ2mito), the protein will be targeted to mitochondria even in the absence of the endogenous SYNJ2BP. Indeed, the overexpression of SYNJ2mito WT in neurons (Fig. S7A) was sufficient to overcome the knock down of SYNJ2BP and relocalized *Pink1* mRNA back onto mitochondria (Fig. 7B and C). This was again specific to *Pink1*, as the localization of *beta actin* transcripts was not altered (Fig. S7B and C). Upon expression of VQL/AAA mutant of the SYNJ2mito construct, the *Pink1* transcript remained cytosolic (Fig. 7B and C). The RNA-binding capacity of SYNJ2 can therefore mediate the localization of *Pink1* mRNA to mitochondria.

Since we have observed that translation is crucial in the *Pink1* mRNA recruitment to mitochondria, we tested whether active translation is necessary for SYNJ2 to localize to the mitochondrial surface. Using a proximity-ligation assay (PLA) SYNJ2 and SYNJ2BP were observed to be closely associated, but upon addition of Puromycin this interaction was reduced to background levels when using mitochondrial proteins outside the RNA adapter complex (PLA of SYNJ2 with RHOT1 and PLA of SYNJ2BP and ATP5b) (Fig. 7D and E). This suggests that in the absence of translation SYNJ2 may be cytosolic, as is the *Pink1* mRNA (compare Fig. 4C), but upon translation of PINK1, SYNJ2 and SYNJ2BP can interact and tether the *Pink1* mRNA to mitochondria.

## Discussion

Our study identifies a neuron-specific mechanism for localizing *Pink1* mRNA to mitochondria and that thereby facilitates the transport of *Pink1* mRNA into axons and dendrites. Application of acute stressors to axons triggers local PINK1-dependent mitophagy (Ashrafi et al., 2014), yet the short half-life of the PINK1 protein makes its transport as a protein into distant regions of the cell highly unlikely. The PINK1/Parkin pathway depends on the ongoing synthesis of PINK1 and is activated by the arrest of PINK1 degradation. How then is the PINK1 protein steadily supplied to distal axons and dendrites? Our finding of a transport mechanism for *Pink1* mRNA and of the dependence of axonal mitophagy on axonal translation establishes a mechanism for local PINK1 translation that can fulfill the requirement for local PINK1 supply.

The failure of local mechanisms for quality control and clearance of mitochondria has been invoked in PD to account for axon degeneration prior to cell death (Cheng et al., 2010; Sliter et al., 2018). Activation of the PINK1/Parkin pathway likely contributes to axonal quality control as it causes acutely depolarized mitochondria to arrest their movement and recruit autophagic and lysosomal markers (Ashrafi et al., 2014; Hsieh et al., 2016). This pathway likely acts in parallel with other forms of mitochondrial clearance from axons. Alternative mechanisms may include the retrograde transport of damaged organelles towards the cell body (Evans and Holzbaur, 2020; Lee et al., 2018; Lin et al., 2017; McWilliams et al., 2018; Miller and Sheetz, 2004). The relative contributions of these mechanisms may vary with the nature of mitochondrial damage and with age; an age-dependent increase in the amount of PINK1- dependent mitophagy occurs in *Drosophila* (Cornelissen et al., 2018) that had not been detected in earlier studies (Lee et al., 2018; McWilliams et al., 2018). Mitochondrial transport has already been known to be crucial for neuronal health by rejuvenating the protein pool of axonal and dendritic mitochondria (Misgeld and Schwarz, 2017). In light of its role in the transport of *Pink1* mRNA and consequently for local mitophagy, mitochondrial transport is therefore important on more than one level for sustaining healthy mitostasis.

Local translation in dendrites is well established (Donnelly et al., 2010; Holt and Schuman, 2013) but axonal translation *in vivo* was controversial. Recent studies, however, demonstrated the presence of translating ribosomes in axons *in vivo*, regardless of the age of the animal (Biever et al., 2020; Ostroff et al., 2019; Shigeoka et al., 2016). All studies report *Pink1* mRNA among the axonally translated transcripts, which is consistent with our findings of *Pink1* transcripts in axons of cultured neurons and optic nerve, whether by qPCR (Fig. 1E and F), *in situ* hybridization (RNAscope, Fig. 2A), or the MS2/PP7-split-Venus approach (Fig. 2D-E). We also detect the local synthesis of a PINK1-N-Kaede construct in axons (Fig 1G and H). Moreover, because inhibition of translation specifically in axons prevented Parkin recruitment to damaged mitochondria (Fig.1D), we have established a functional significance for its axonal translation.

Axonal transcripts have been found to regulate axon growth, maintenance and regeneration after injury (Andreassi et al., 2010; Ben-Yaakov et al., 2012; Hanz et al., 2003; Hengst et al., 2009; Sotelo-Silveira et al., 2008; Terenzio et al., 2018; Verma et al., 2005; Yoon et al., 2012). In addition, mitochondria have come into the spotlight as centers for axonal translation (Cioni et al., 2019; Cosker et al., 2016; Spillane et al., 2013). However, to the extent these transcripts have been localized and their transport mechanisms examined, these transcripts are thought to travel as independent RNP granules. *Pink1* mRNA, however, did not form such separate RNP granules. Instead, *Pink1* mRNA consistently colocalized with mitochondria and thus offers a different model for a means of transport. We observed all processive movements of *Pink1* mRNA to be coupled with mitochondrial movements (Fig. 3C) and measures that are known to alter mitochondrial motility, like the overexpression of kinase-active PINK1 protein, were reflected in the motility of the *Pink1* mRNA (Fig. 3D). When transcripts were observed without a detectable associated mitochondrion, they were either stationary or engaged in only short-range and often erratic movements (Fig. 3B). Sensitive detection of *Pink1* mRNA by RNAscope found the transcript on almost all mitochondria in axons and dendrites (Fig. 2A and B), which would include those that were stationary or moving retrograde before fixation. Thus, the association of transcript and mitochondrion is probably not solely a mechanism for delivery from the cell body to the periphery, but may also facilitate the constant translation of PINK1 as is necessitated by its role in mitochondrial quality control (Narendra et al., 2010).

The localization of the transcript to neuronal mitochondria was verified using both RNAscope and a live imaging approach using split-Venus fluorophore reconstitution on a stem/loop tagged *Pink1* construct. RNAscope has the clear advantage of detecting the endogenous transcript, and was more easily applied to distal axons. However, it was not appropriate for live imaging of mRNA transcript. For this, we turned to the MS2/PP7-split-Venus method (Wu et al., 2014). Although it easily visualized transcripts in the soma and dendrites, detection of the *Pink1* mRNA was challenging in axons. Split fluorophore reconstitution prevents detectable background from the unassembled components. It adds, however, to the complexity of the system, as the loss of any one of the ternary complex partners during transport leads to loss of signal. Genomic integration of the stem loops, as was done for *beta actin* in a transgenic mouse model (Lionnet et al., 2011), would permit imaging of endogenous transcripts, but would still face the need for two well-transported fluorophore constructs (compare Fig S2C). Transient overexpression, nevertheless, allowed the efficient screening of constructs in order to find *cis*-acting sequences in the *Pink1* mRNA (Figure 4). Because the *Pink1* mRNA was mitochondrial in all neuronal compartments, we were able to use transport in dendrites and mitochondrial colocalization in somata for defining how *Pink1* mRNA localized to mitochondria and entered neuronal processes.

We examined the mechanism of the association between *Pink1* mRNA and mitochondria and their co-transport by determining both the necessary *cis*-elements and requisite *trans*-acting factors. We determined that there are at least two sequences required for the mitochondrial localization of the transcript. Deletion of the start codon or inhibition of translation by puromycin indicated that translation of the PINK1 protein is one requirement for mitochondrial localization of the transcript (Fig. 4C and D). The N-terminal MTS of PINK1 may interact with the receptors of the TOM complex as soon as the nascent chain emerges from the ribosome and thereby confer mitochondrial localization to the transcript. But, though translation of PINK1 is necessary, translation of the MTS is not sufficient: a transcript in which this MTS properly directed its BFP-tagged translation product into mitochondria (*Pink1 5’UTR+MTS-BFP*, Fig. 4E) remained cytosolic. Therefore, at least one additional sequence must be necessary and a comparison of the constructs in Figure 4 implies that it is likely found between base pairs 103 and 625 of the *Pink1* open reading frame (compare *Pink1 5’UTR+N-BFP* and *Pink1 5’UTR+MTS BFP*, Fig. 4A and B). If the *Pink1 5UTR’+MTS-BFP* transcript is transiently associated with the mitochondrion while BFP is being translated and translocated into the matrix, it does not remain associated with mitochondria as does full length *Pink1* mRNA. In yeast, interaction between the nascent chain and mitochondria mediates the co-translational targeting of the *Atp2p* mRNA to the mitochondrial surface (Garcia et al., 2010), but there is an additional role of its 3’UTR for its mitochondrial localization (Margeot et al., 2002). This dual requirement is reminiscent of the bipartite targeting signal of *Pink1* mRNA.

In exploring the *trans*-acting factors required in neurons, we discovered that knock down of the mitochondrial outer membrane protein SYNJ2BP is sufficient to remove *Pink1* mRNA from mitochondria (Fig. 5A and B). Once released from mitochondria, *Pink1* mRNA was sequestered in cytoplasmic granules containing the RNA helicase DDX6 (Fig. 5C and S5B). This sequestration did not represent a uniform change in cellular mRNA; *beta actin* mRNA distribution was unaltered (Fig. S5C and D). Thus the relocalization of the *Pink1* transcript was likely to be a direct consequence of its inability to associate with mitochondria. SYNJ2BP itself has been suggested to have RNA-binding capabilities (Qin et al., 2020), but our observation that the mitochondrial localization of the *Pink1* transcript was neuron-specific led us to explore the expression patterns of its interactors. Unlike the SYNJ2BP-interacting ER protein RRBP1 (Hung et al., 2017), which was much more abundant in fibroblasts (Fig. 6C), the mitochondria-specific splice-variant of SYNJ2, SYNJ2a, was five-fold higher expressed in neurons over fibroblasts (Fig. 6B). Overexpression of Synj2a and SYNJ2BP was sufficient to target *Pink1* mRNA to mitochondria in COS-7 cells (Fig6 D and E). PINK1 has been reported to alter proteins important for mitochondrial dynamics and ultrastructure (Chen and Dorn, 2013; Tsai et al., 2018; Wang et al., 2011); altered PINK1 biogenesis may therefore account for the clustering of mitochondria seen by us and others upon Synj2a overexpression (Nemoto and De Camilli, 1999). Alternatively the clustering may arise from other, yet to be identified substrates of the SYNJ2a/SYNJ2BP RNA tethering complex, or lipid alterations due to the inositol 5’-phosphatase activity of SYNJ2.

SYNJ2 is best known for its inositol 5’-phosphatase function during endocytosis, but we demonstrated an RNA-binding capacity of SYNJ2 (Fig. 7A) and showed that the engineered presence of this protein on the mitochondrial surface was sufficient to localize *Pink1* mRNA at mitochondria, even in the absence of SYNJ2BP (Fig. 7B and C). Thus the only essential function of SYNJ2BP for the localization of the mRNA is its ability to recruit SYNJ2. Mutations that abolished RNA binding by the RNA-Recognition Motif in SYNJ2 demonstrated that it is indeed this capacity of SYNJ2 that is needed. It is interesting to speculate whether mitochondrially localized SYNJ2a would still be able to interact with its endosomal binding partners, thereby forming a mito-Endosome contact site. Fittingly, local translation in axons has been reported to occur at such interfaces (Cioni et al., 2019).

The mRNA tether we propose would thus entail SYNJ2BP anchored in the outer membrane and associating with its partner SYNJ2, which in turn holds the mRNA. SYNJ2a and SYNJ2BP were shown to directly interact in yeast two-hybrid screens and in purified proteins by virtue of a PDZ domain in SYNJ2BP and an interacting motif in SYNJ2a (Nemoto and De Camilli, 1999). We have confirmed that the proteins are associated *in situ* by PLA, however the association was greatly diminished when translation was inhibited (Fig. 7E). This translation-dependence is likely significant for the translation-dependent localization of *Pink1* mRNA to neuronal mitochondria. Thus localization appears to involve two mechanisms acting in parallel and reinforcing one another. One model that combines these components would first envision active translation of the PINK1 MTS leading to co-translational protein import of the nascent chain and hence the relocation of the ribosome and associated *Pink1* mRNA to the mitochondrial surface. Thereafter, the interaction of the mRNA and the mitochondrion is further stabilized through the SYNJ2/SYNJ2BP tether. This mechanism allows the *Pink1* transcript to move together with mitochondria in neurons while providing a constant local source for freshly synthesized PINK1 protein.

Our observation also raises the question as to whether the predicted RRM domains of all Synaptojanins are also capable of RNA-binding. *Beta actin* mRNA was not redistributed upon expression of SYNJ2mito (Fig. S7B and C) and SYNJ1, while containing an RRM, could not replace SYNJ2 for localization of *Pink1* mRNA (Fig. 6D and E). Thus, there is a substrate specificity to the RNA-binding capacity of the synaptojanins.

It will now be critical to determine what other mRNAs may be bound by SYNJ2. Mitochondrial transcripts are among the most abundant RNAs found in axons, and at least some mitochondrial proteins have a comparatively short half-life that may require their local translation to ensure mitostasis in axons (Harbauer, 2017; Misgeld and Schwarz, 2017). Both *Atp5f1b* and *Cox4i* transcripts were enriched on neuronal mitochondria (Fig. S2E and F), as well as *Cox7c* identified in the accompanying manuscript (Cohen et al., 2021), which makes them likely candidates for binding to SYNJ2a. Although loss of *Pink1* does not compromise the viability of cultured neurons (Kitada et al., 2007), neuronal health deteriorated upon knock down of SYNJ2BP; this effect may indicate that there are other transcripts, vital to neurons, that depend on this tether for localization and transport.

PD primarily affects neurons. Our observations that mitochondrial localization of the *Pink1* mRNA may be specific to neuronal cells (Fig. 6A) and that local translation and mitochondrial tethering through SYNJ2BP is required for the full activation of the PINK1/Parkin pathway in axons (Fig. 1D and 5E) point to the exceptional challenges that are faced by neurons as they seek to maintain healthy mitochondria in distal regions of their arbors. Effective transport of mRNA for mitochondrial proteins, whether on moving mitochondria or as RNP granules, likely contributes substantially to preserving neurites from degeneration.

## Acknowledgments

We are grateful for plasmids shared with us by P. de Camilli (Yale University; FLAG-SYNJ2a and SYNJ1-FLAG). We thank L. Mkhitaryan and J. Lindner for technical support and the members of the T.L. Schwarz laboratory for their support and many fruitful discussions. We are grateful to S. Vasquez and M. Sahin’s laboratory for assistance with primary hippocampal cultures; D. Tom, L. Ding, M. Ocaña and A. Begue from Harvard NeuroDiscovery Center’s/Neurobiology Imaging Facility Enhanced Neuroimaging Core (NINDS P30 Core Center grant no. NS072030), as well as R. Kasper and M. Spitaler from the Imaging facilities of the MPIs for Neurobiology and Biochemistry, respectively, for assistance with live-cell imaging. This work was supported by grants from the National Institute of Health R01NS107490 and R01GM069808 to TLS, and P30HD018655 to the IDDRC Cellular Imaging and Molecular Genetics Cores. ABH was supported by a Howard Hughes Medical Institute fellowship from the Jane Coffin Childs Memorial Fund for Medical Research. Since 2019, she is an independent research group leader and a Rudolf-Mößbauer tenure track professor at Technical University of Munich Institute for Advanced Study and the Institute of Neuronal Cell Biology. In this role she is supported by the Max Planck Society and the Deutsche Forschungsgemeinschaft (DFG, German Research Foundation, HA 7728/2-1 and under Germany’s Excellence Strategy within the framework of the Munich Cluster for Systems Neurology (EXC 2145 SyNergy – ID 390857198).

## Author Contributions

ABH and TLS conceived of the project and wrote the manuscript. ABH designed and conducted most of the experiments. SW was responsible for RNAscope and superresolution imaging and the PINK1-Kaede imaging. JTH was responsible for imaging Atp5f1b and Cox4i live cell imaging and PLA experiments. WG contributed the analysis of PINK1 stability in human iPSC-derived neurons. GA conducted initial experiments for Parkin translocation in the presence of CHX, MO conducted qPCR experiments and ZC assisted with COS cell experiments and cloning. RC, CW and ZH were responsible for expression of PINK1 constructs in mouse retinal ganglion cells.

## Figure Legends

**Figure S1.**
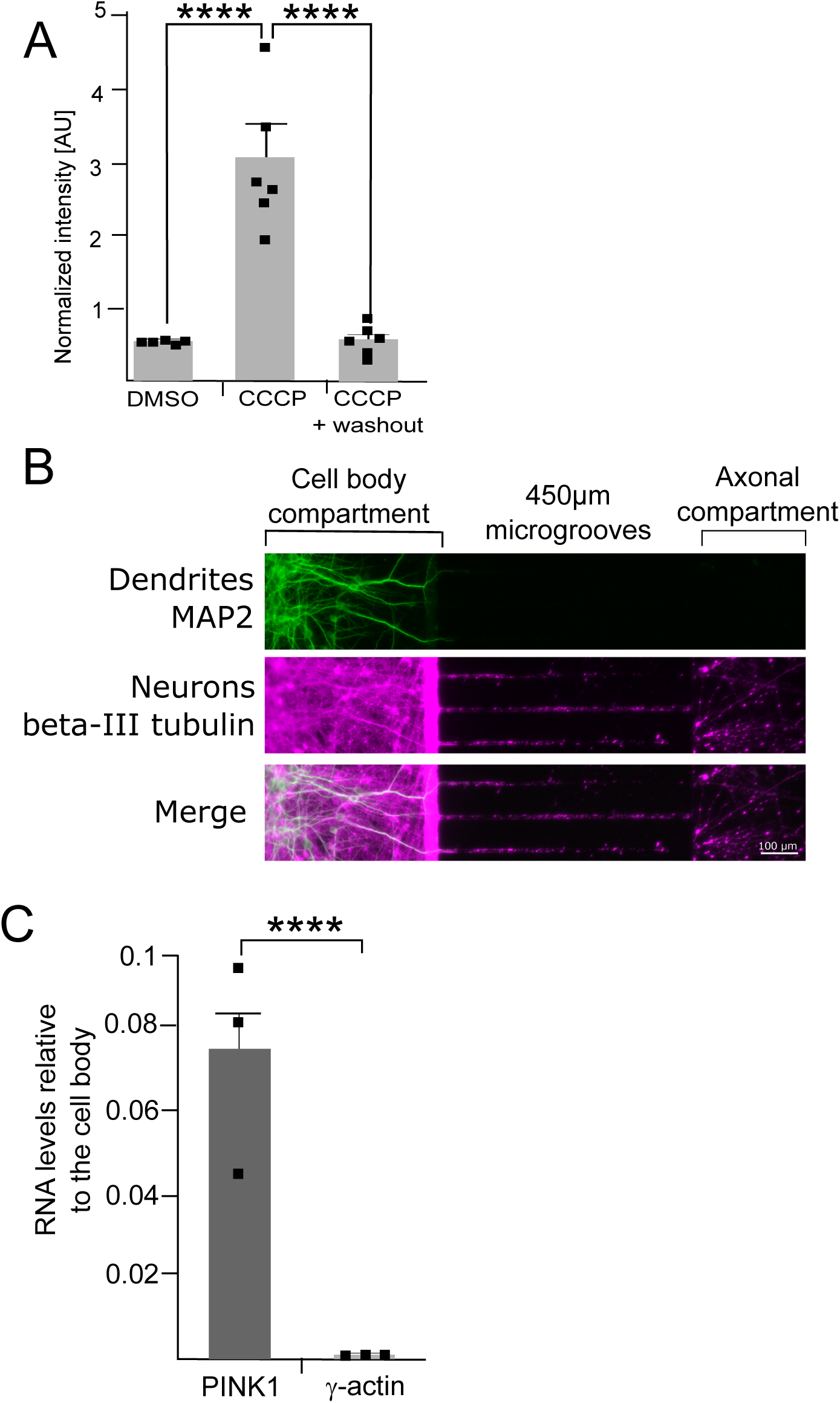
corresponding to Figure 1. Microfluidic chambers exclude dendrites and lack detectable cell body transcript. (A) Quantification of PINK1 stabilization in human iPSC- dervied cortical neurons upon CCCP treatment (4.5h 20µM) and 30 min after washout of CCCP. Intensity is shown normalized to GAPDH signal. Data is shown as mean ± SEM; ANOVA with Tukey’s multiple comparisons test, n=6 biological repeats per condition (B) Immunolabeling of neurons grown in a microfluidic chamber for a dendritic (MAP2) and a neuronal (beta-III tubulin) marker. (C) qPCR analysis of RNA levels in the cell body and axonal compartment. Please note that the total amount of RNA isolated from the axonal side is approximately 20 times lower than the cell body. The same data normalized to mitochondrial rRNA is presented in Figure 1E.

**Figure S2.**
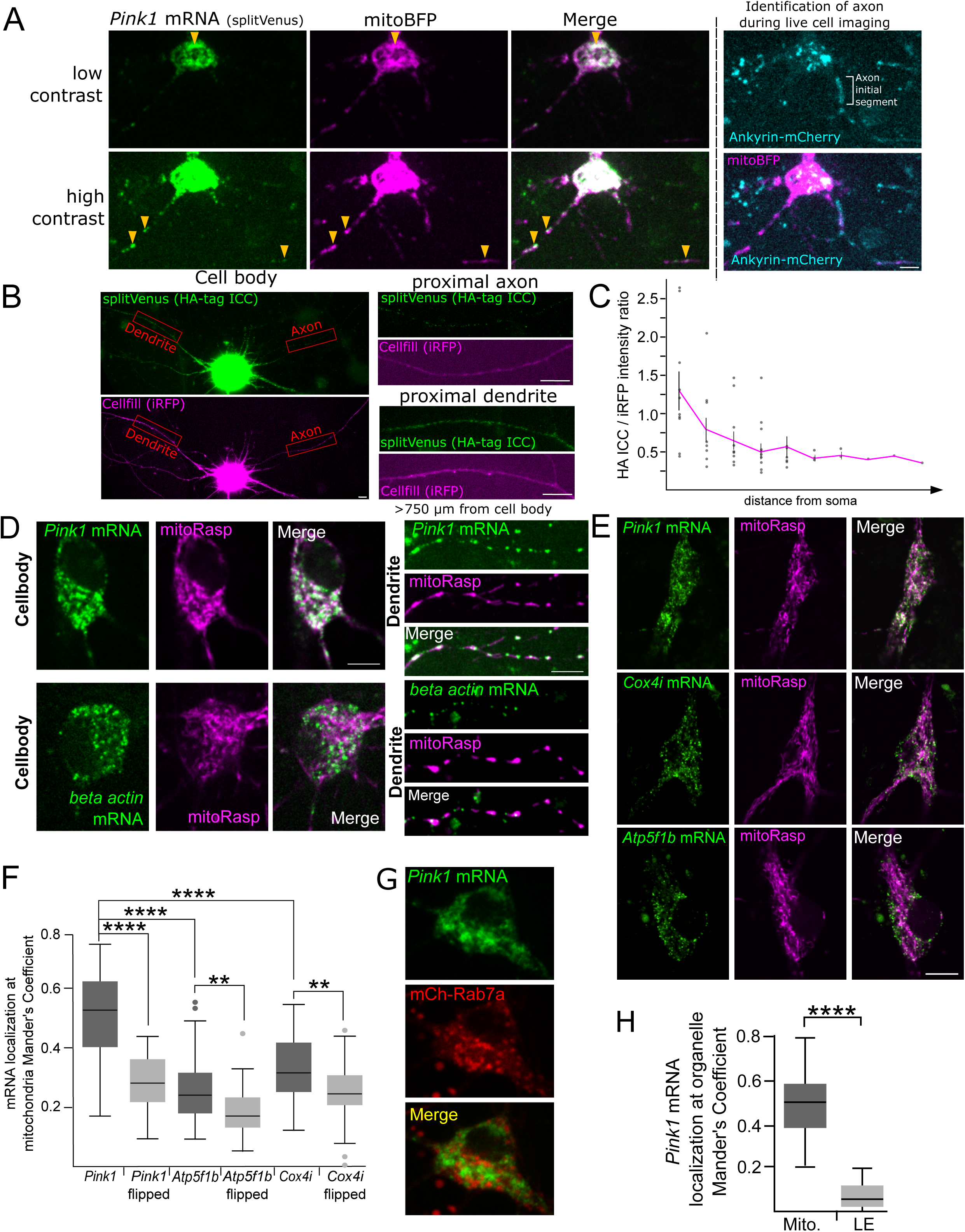
corresponding to Fig 2. Analysis of the *Pink1, Cox4i and Atp5f1b* mRNA localization using the MS2/PP7/-split-Venus method. (A) Representative image for kinase-dead *Pink1* mRNA localization at mitochondria in all neuronal compartments using MS2/PP7- split-Venus to visualize *Pink1* transcripts, mitochondrially targeted BFP (mitoBFP) to visualize mitochondria and AnkyrinG-mCherry to identify the axon. (B and C) Representative image and quantification of the splitVenus subunits detected by their HA-tag and an iRFP cell-fill. Axons were traced as many fields of view as possible using the iRFP cell-fill and the mean intensity after background correction was used to calculate the ratios presented in (C). The magenta line displays the mean ratio for each field of view (approx. 500 µm) and error bars correspond to SEM. (D) Live cell imaging of *Pink1* and *beta actin* RNA in hippocampal neuron cell bodies and dendrites and as well as a mitochondrial marker (mRaspberry targeted to the mitochondrial matrix, mitoRasp). (E) Representative images of *Pink1*, *Cox4i* and *Atp5f1b* transcripts visualized with the MS2/PP7-split-Venus method and mitochondria detected by mitochondrially targeted Raspberry (mitoRasp). (F) Mander’s correlation coefficient between the indicated transcripts and mitochondria. Student’s t-test, n = 29-31 cell bodies, p<0.01(**), p<0.0001 (****). (G) Representative images for kinase-dead *Pink1* transcript localization in neuronal cell bodies relative to late endosomes labelled with mCherry(mCh)-Rab7a and mitochondria labelled with mitoBFP. Scale bar = 10 µm. (H) Quantification with Mander’s correlation coefficients of the extent of transcript colocalization in (G) with mitochondria (Mito.) and with Late Endosomes (LE). Student’s t-test, n = 20 cell bodies; p<0.01 (**), p<0.0001 (****). Scale bars = 10 µm (except (A)).

**Figure S3.**
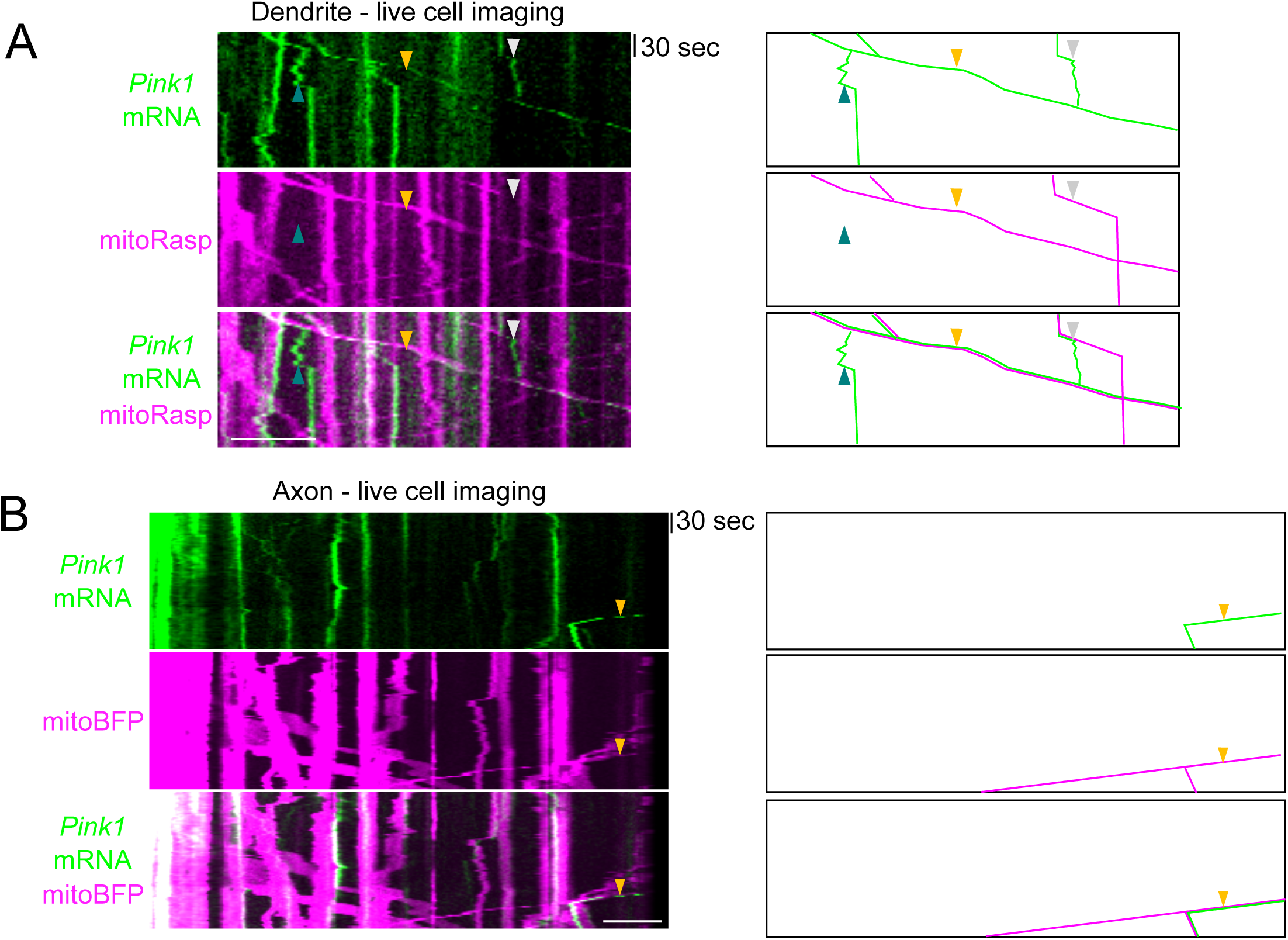
corresponding to Figure 3. *Pink1* mRNA is transported with mitochondria in axons and dendrites. (A) Repetition of Figure 3B alongside a schematic that represents a likely interpretation of selected elements in the kymograph. Co-transport of mRNA and mitochondria is present (yellow arrowhead). At the grey arrowhead, a *Pink1* mRNA particle appears to dissociate from a moving mitochondrion and, after a period, to reassociate with a different moving mitochondrion. Another potential dissociation and brief episode of short-range mitochondrially independent movement of a *Pink1* mRNA particle is marked with a blue arrow head. Scale bar = 10 µm. (B) Kymograph and schematic of *Pink1* mRNA and mitochondrial co-transport (yellow arrowhead) in a proximal axon. Scale bar = 10 µm

**Figure S4.**
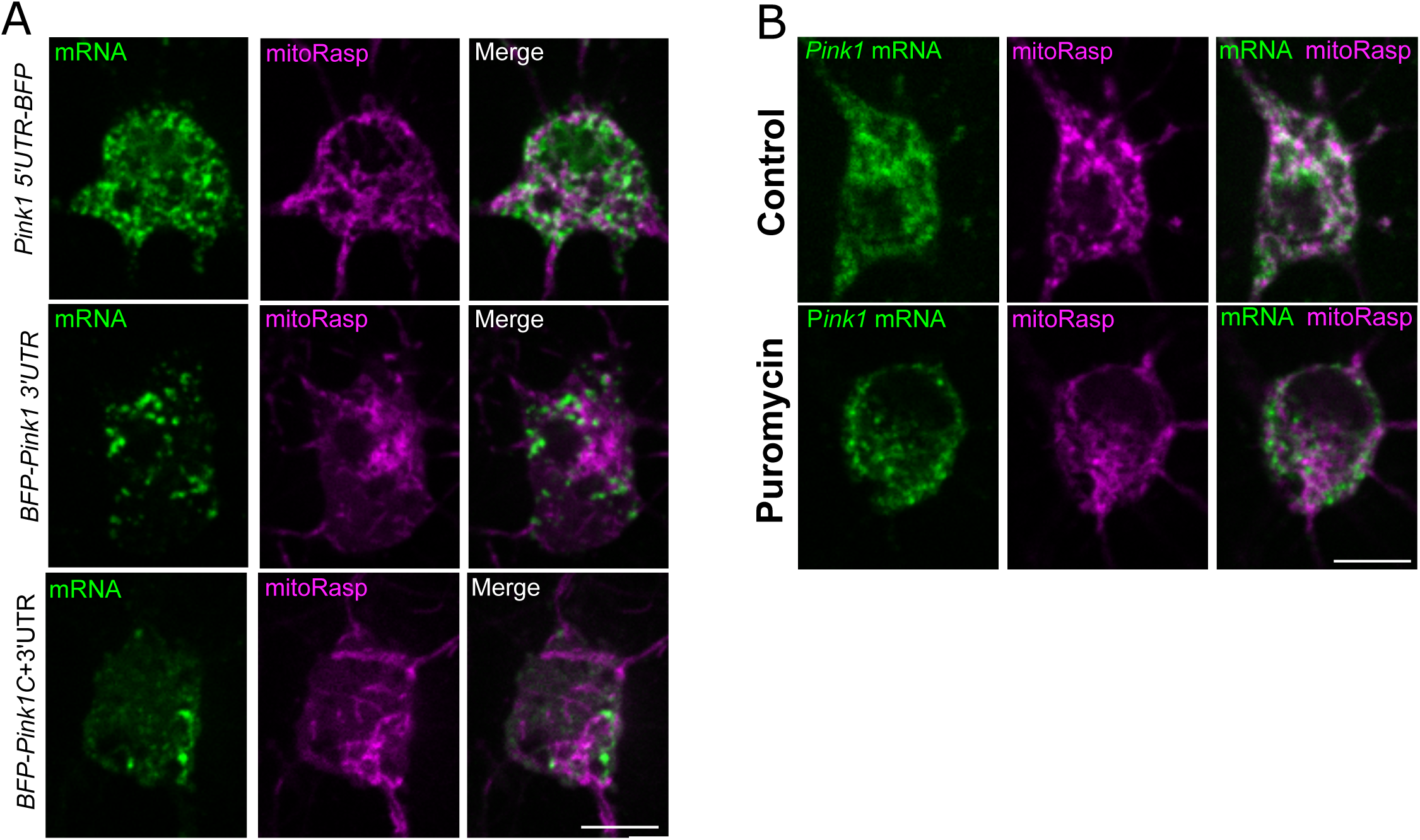
corresponding to Figure 4. Representative images of Pink1-BFP. (A) Representative images for *Pink1 5’UTR-BFP, BFP-Pink1 3’UTR* and *BFP-Pink1 C+3’UTR* transcript localization. (B) Representative localization of full length *Pink1* (kinase dead) transcript in the presence or absence of puromycin. Scale bars = 10 µm.

**Figure S5.**
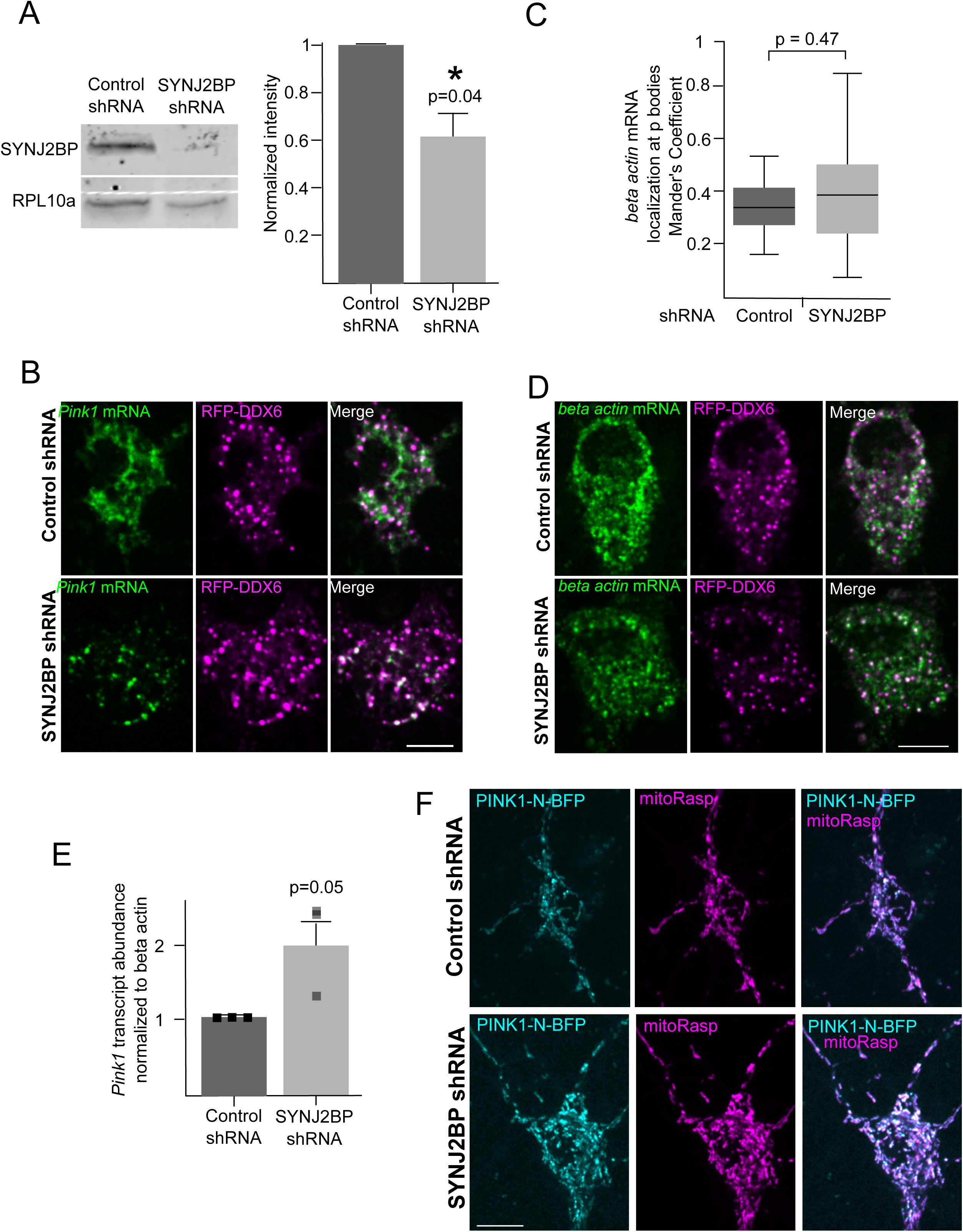
corresponding to Figure 5. Knock down of SYNJ2BP does not alter *beta actin* mRNA localization nor PINK1 translation or RNA stability. (A) Representative Western Blot and quantification of SYNJ2BP protein levels in neurons treated with control or SYNJ2BP shRNA for three days. Band intensity was normalized to Rpl10a and control shRNA. Data is shown as mean ± SEM; Student’s t-test; n = 3 experiments, p<0.05 (*). (B) Representative images of neuronal cell bodies expressing the split-Venus reporter for the *Pink1* transcript and RFP-tagged DDX6. (C) Colocalization as analyzed with Mander’s correlation coefficient between the *beta actin* transcript and P-bodies marked with RFP-DDX in control and SYNJ2BP shRNA treated neuron cell bodies. Student’s t-test, n = 16-17 cell bodies. (D) Representative images from control or SYNJ2BP shRNA treated neuron cell bodies illustrating the localization of *beta actin* mRNA with respect to DDX6. (E) Pink1 transcript abundance was analyzed by qPCR in control of SYNJ2BP shRNA treated neurons. Data is presented as mean ± SEM; Student’s t-test, n=3 biological repeats per condition (F) Representative micrographs of PINK1-N-BFP (compare Figure 4) expression in control and SYNJ2BP knock down neurons. Scale bar = 10 µm.

**Figure S6.**
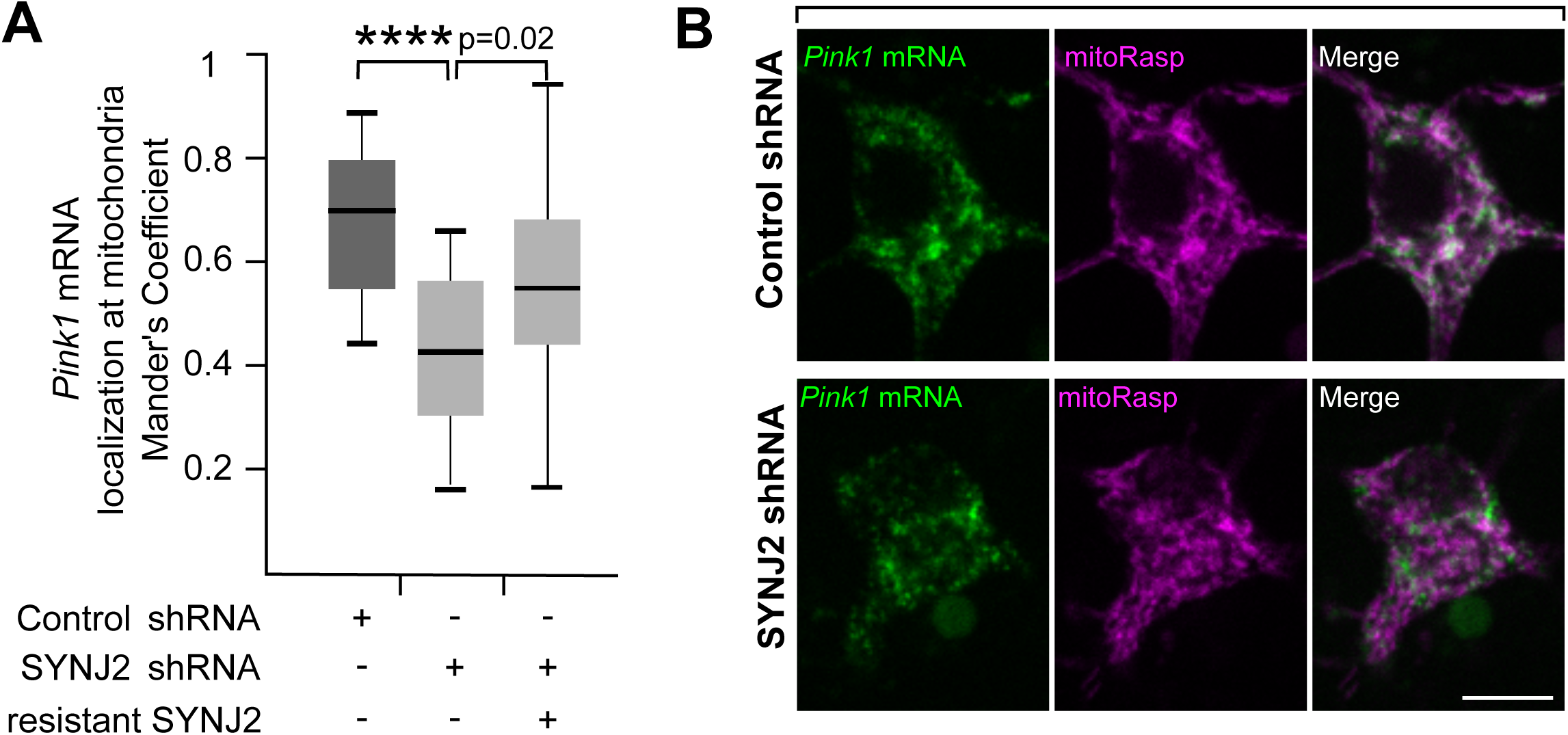
corresponding to Figure 6. Knock down of SYNJ2 affects *Pink1* mitochondrial localization in neurons. Quantification (A) and representative images (B) of mitochondrial localization of the *Pink1* transcript visualized by the split-Venus method and mitoRasp in control and SYNJ2 shRNA treated hippocampal cell bodies. Expression of an shRNA-resistant SYNJ2 restored localization of Pink1 transcripts to mitochondria. Student’s t-test, n = 14-20 cell bodies, p<0.05 (*) and p<0.0001 (****). Scale bar = 10 µm.

**Figure S7.**
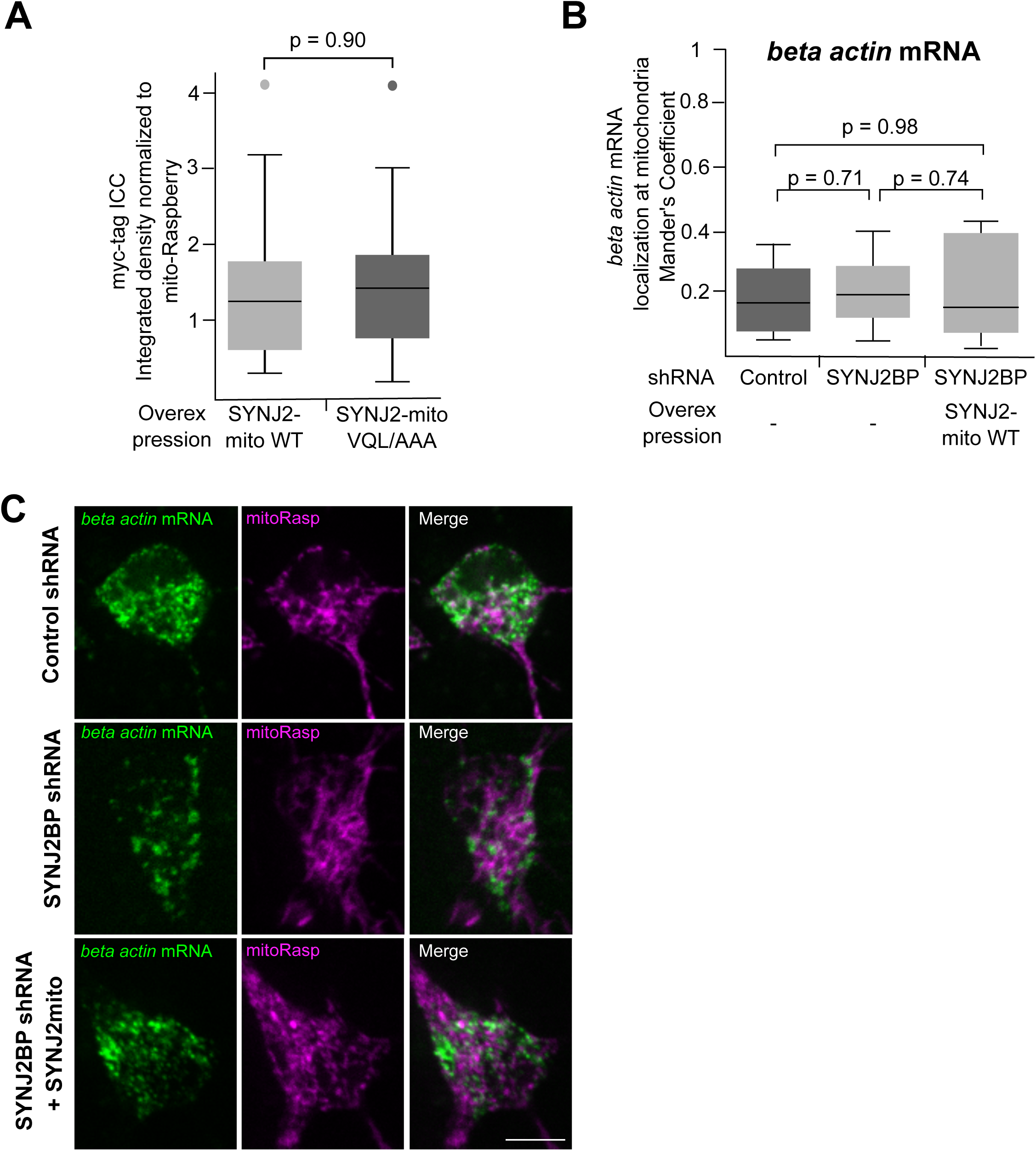
corresponding to Figure 7. Expression of SYNJ2mito does not alter *beta actin* transcript localization. (A) SYNJ2-mito VQL/AAA is similarily expressed as its WT counterpart as analyzed by immunocytochemistry (ICC) against its myc-tag normalized to the co-transfected mitoRaspberry. Student’s t-test, n = 20-21 neurons. (B) Colocalization as quantified with Mander’s correlation coefficient between mitochondria and *beta actin* transcripts. (C) Representative images from control or SYNJ2BP shRNA treated neuron cell bodies with and without expression of SYNJ2mito. Student’s t-test, n = 11-18 cell bodies; Scale bar = 10 µm.

**Supplemental movie 1 corresponding to Figure 3.** *Pink1* mRNA is transported with mitochondria in dendrites.

**Supplemental movie 2 corresponding to Figure 3.** This movie illustrates possible association and dissociation events of *Pink1* mRNA and mitochondria in a dendrite. The movie is played twice. In the first part a white arrow indicates a stationary *Pink1* mRNA particle (arrow) that associates with a moving *Pink1* mRNA on a moving mitochondrion (arrowheads). The second part illustrates a moving *Pink1* mRNA particle that seems to split and while one part keeps moving on (arrowhead) the other stays behind (arrow). There is no detectable mitochondrial signal at the position of the stationary *Pink1* mRNA particle.

**Supplemental movie 3 corresponding to Figure 3.** *Pink1* mRNA is transported with mitochondria in axons. This movie demonstrates a detail of the kymograph presented in Fig. S2. At 18’ a mitochondrion-associated mRNA particle (marked with arrowheads) enters from the right, then pauses. At 44’, a second mitochondrion and its associated mRNA, which had been stable (arrowheads), begins to move.

## Supplemental information

### Key Resources Table

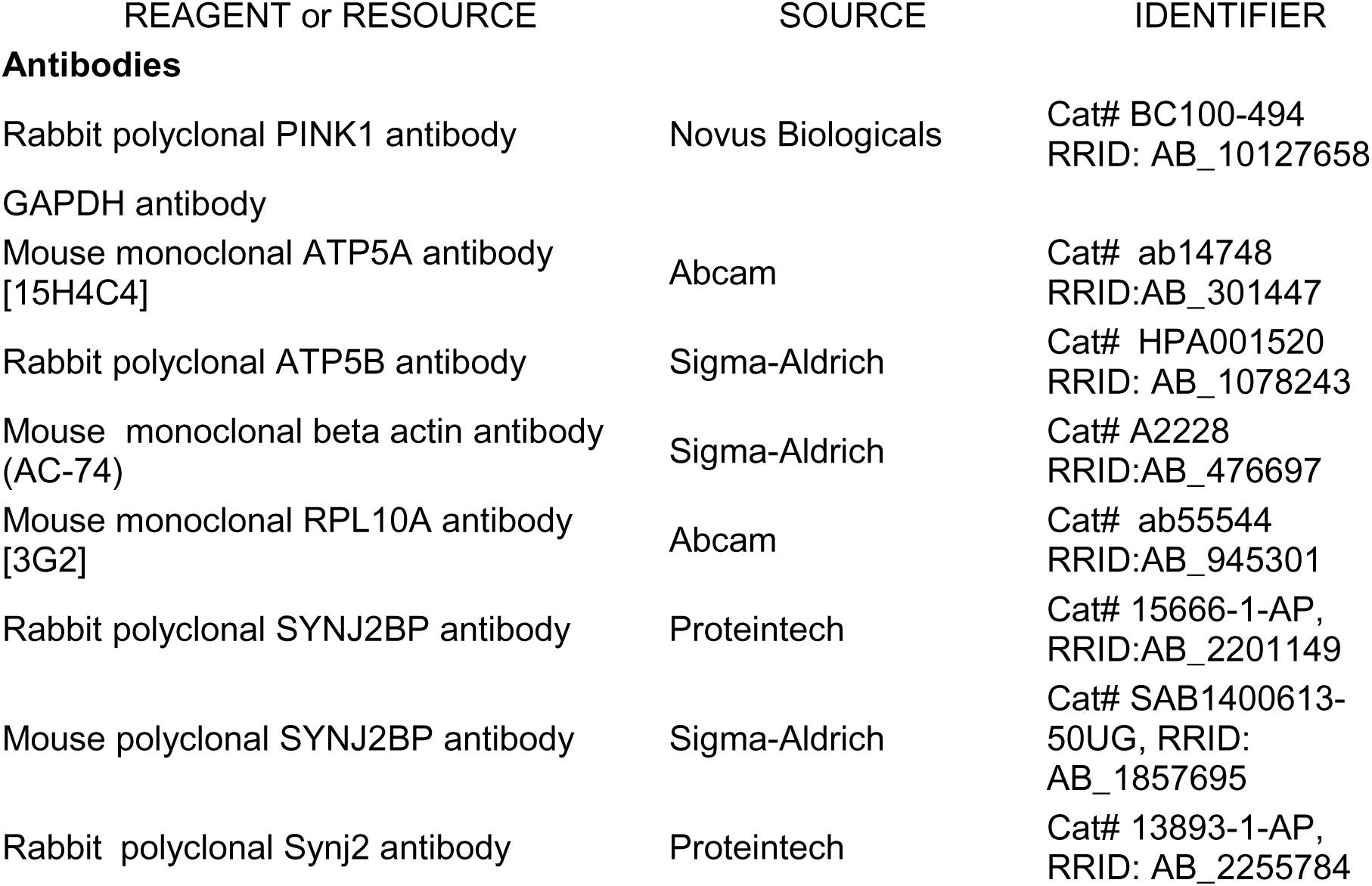
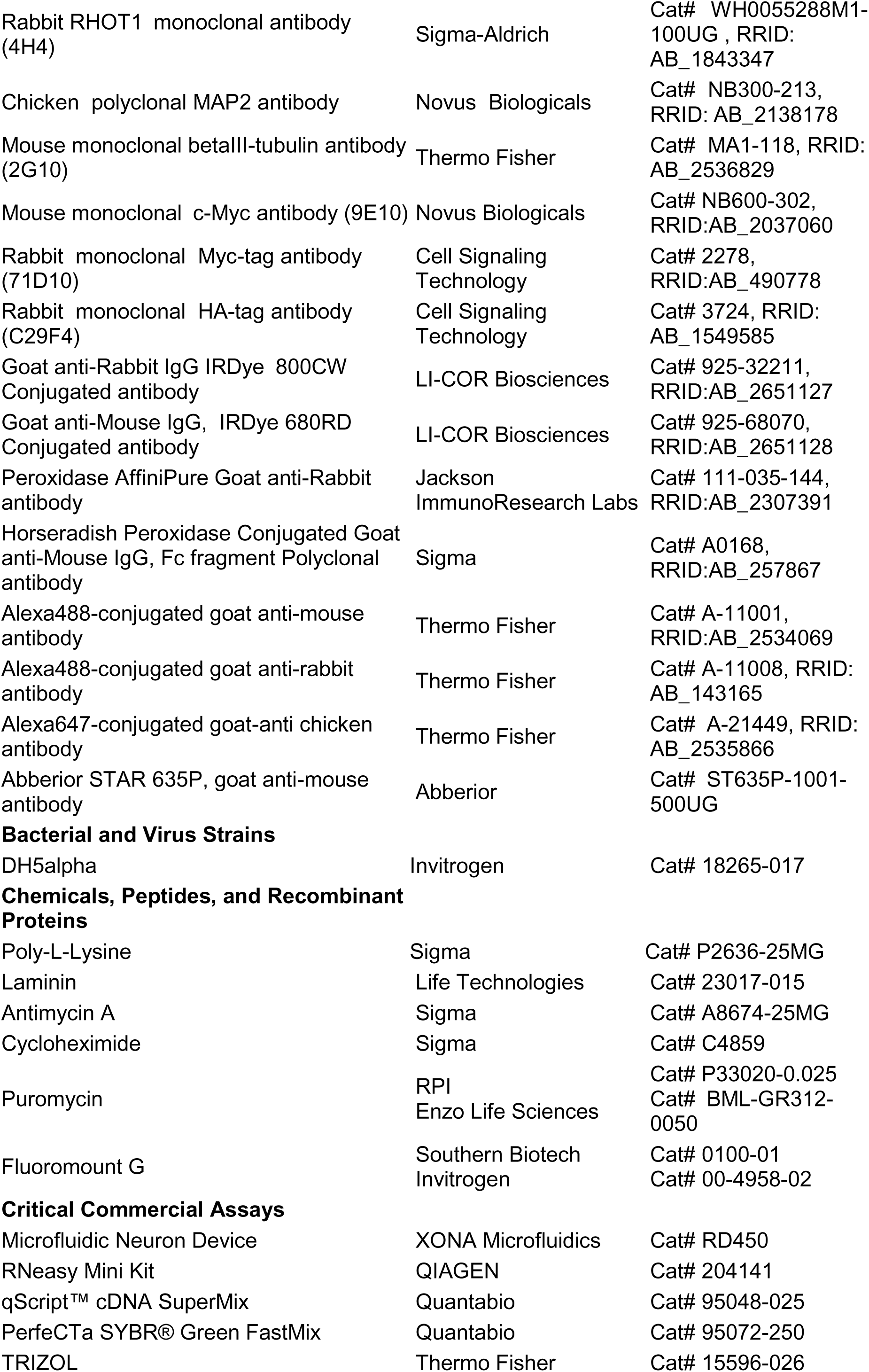
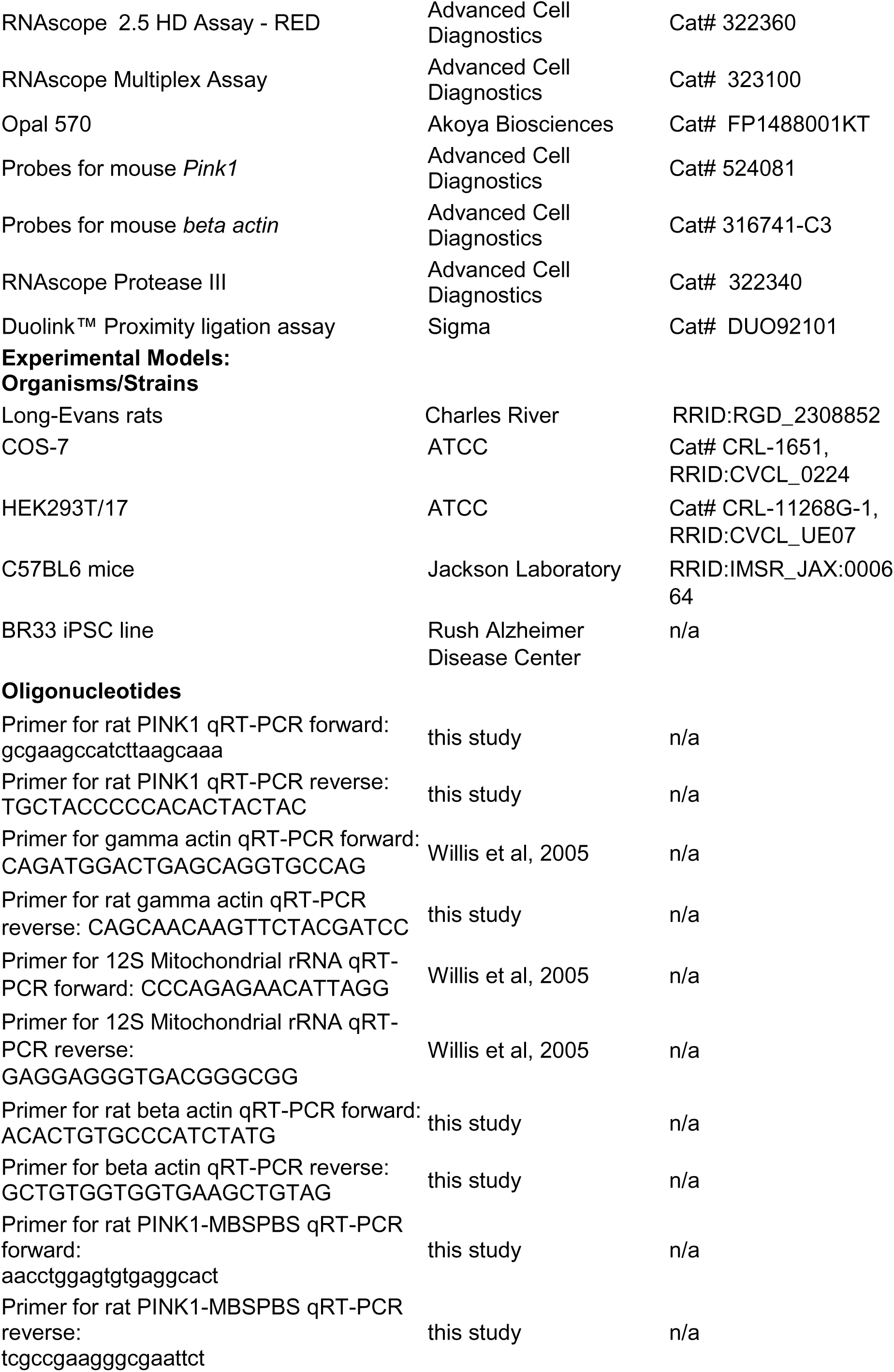
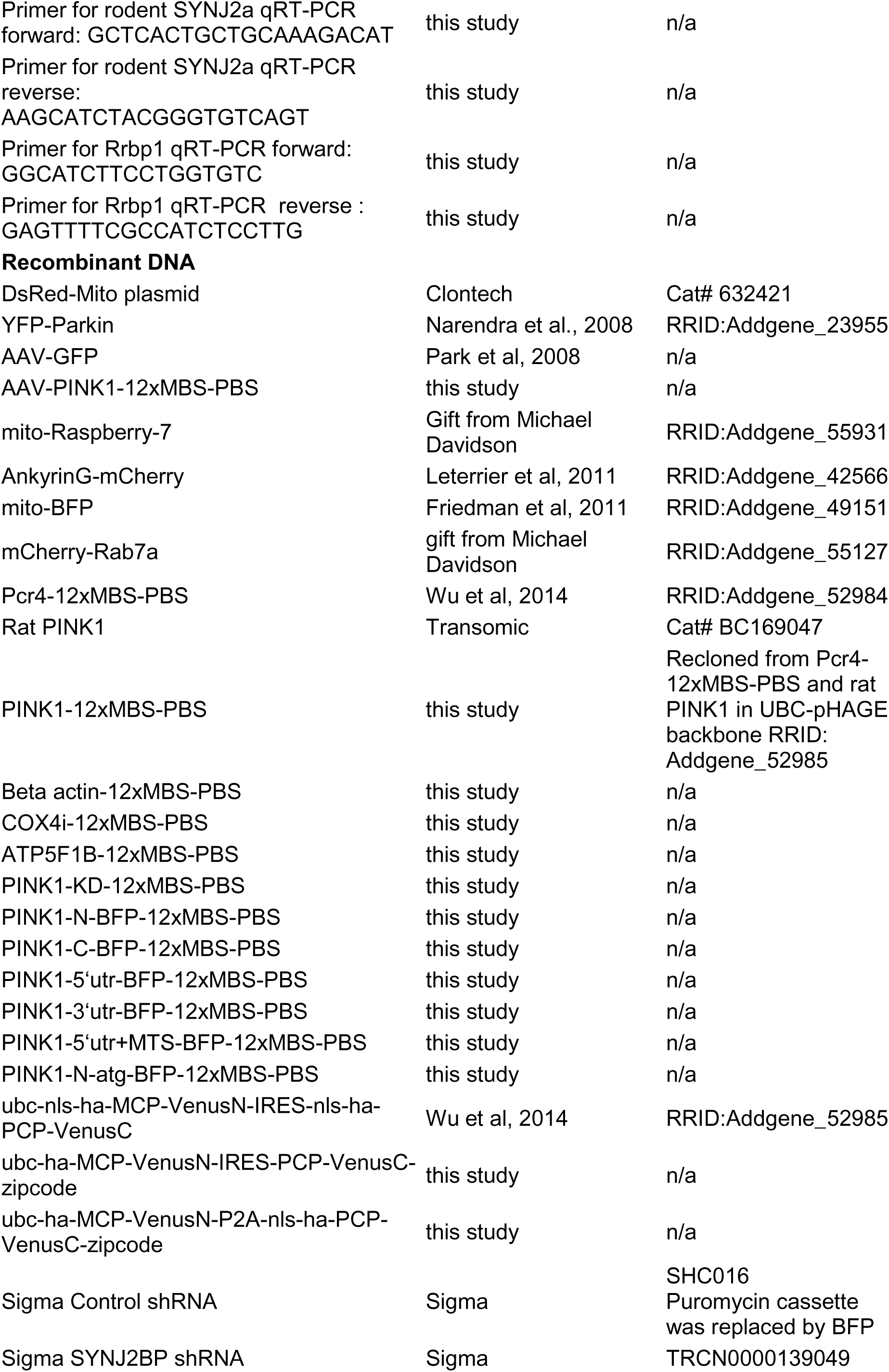
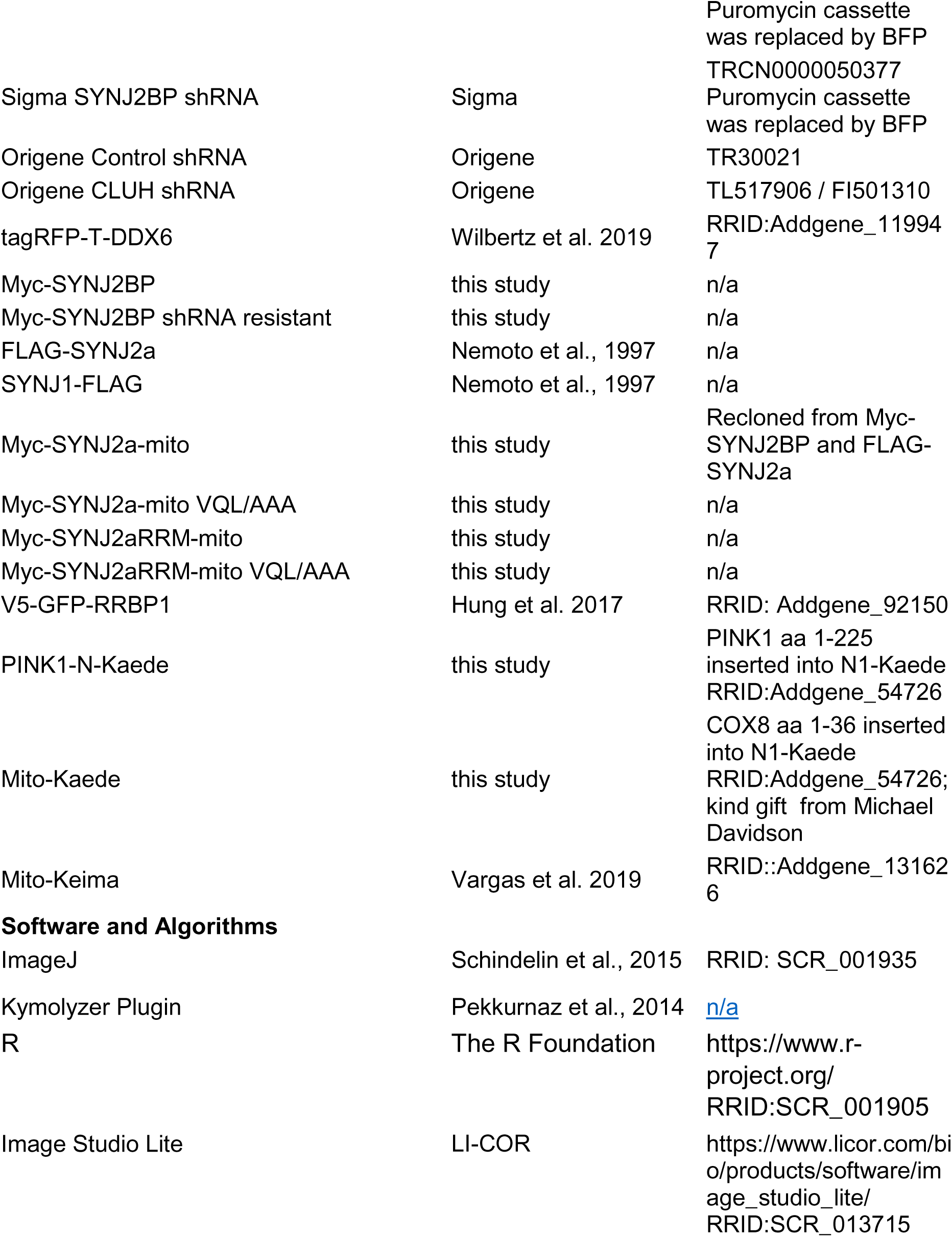

### Contact for Reagent and Resource Sharing

Further information and requests for resources and reagents should be directed to and will be fulfilled by the corresponding author Thomas L. Schwarz.

### Experimental Model and Subject Details

Primary fibroblast and neuron cultures were obtained from E18 (embryonic day 18) Long/Evans rat pups or on E16.5 from mouse pups. Pregnant females from timed matings were delivered from Charles River Laboratories and housed overnight in the animal facility. Rat and mouse procedures were approved by the Institutional Animal Care Committee (IACUC) at the Boston Children’s Hospital and the MPI of neurobiology. BR33 iPSCs were obtained from Rush Alzheimer Disease Center.

### Cell culture preparation

Primary cell cultures were prepared as described in Shlevkov et al. (2016). Hippocampal neurons were obtained by euthanizing the pregnant female with CO_2_ and recovery of the E18 embryos from the abdomen. Hippocampi were dissected and placed in chilled dissociation medium (Ca2+-free HBSS with 100 mM MgCl2, 10 mM kynurenic acid, and 100 mM Hepes), and enzymatically dissociated with Papain/l-cysteine (Worthington Biochemical Corporation). After addition of Trypsin inhibitor (Sigma-Aldrich), tissue was triturated 10-15 times with a P1000 pipet until clumps disappeared. Neurons were resuspended in Neurobasal medium supplemented with B27 (Gibco/Life Technologies), L-glutamine, and penicillin/streptomycin (NB+PSG+B27) and plated on 20 μg/mL poly-L-Lysine (Sigma-Aldrich) and 3.5 μg/mL laminin (ThermoFisher Scientific) coated glass bottom plates (CellVis) or acid washed glass coverslips (1.5mm, Warner Instruments). 50% of the medium was replaced every 2-3 days with fresh NB+PSG+B27. Transfections were performed at day *in vitro* (DIV) 5-7 and imaging at DIV7– DIV9. Fibroblasts were obtained from E18 rat embryos by standard methods and cultured in DMEM+20% FBS. Cells were maintained in T75 flasks or frozen in 10% DMSO for future use.

iPSCs were plated on Matrigel-coated plates (Corning, 354234) and cultured growth factor reduced mTeSR media (StemCell Technologies, 05857) supplemented with ROCK inhibitor (10 uM; StemCell Technologies #72304) at a density of 100K cells/cm^2^. iPSCs were then transduced with lentivirus packaged pTet-O-NGN2-puro and Fudelta GW-rtTA plasmids. NGN2-transduced iPSCs were thawed in StemFlex media with ROCK inhibitor. Once cells were reached 75-80% confluency (day 1), cells were exposed to KnockOut media (Gibco 10829.018) supplemented with KnockOut Serum Replacement (Invitrogen 10928-028), 1% MEM non-essential amino acids (Invitrogen 11140), 1% GlutaMAX (Gibco 35050061) and 0.1% BME (Invitrogen 21985-023) (KSR) containing doxycycline (2 μg/ml, Sigma, D9891-5g) to induce NGN2 expression. On day 2, the media was changed to equal volumes of KSR and N2B media (DMEM F12 supplemented with 1% GlutaMAX, 3% dextrose and N2-Supplement B; StemCell Technologies 07156) with puromycin (5 μg/ml; Life Technologies, A11138-03) and doxycycline to select for transduced cells. On day 3, cells were feed with N2B media containing media with B27 (1:100; Life technologies, 17504-044), puromycin, and doxycycline. On day 4, the cells were dissociated with Accutase (Gibco, A11105) and frozen down in freezing media containing 1:1 ratio of 20% DMSO and Neurobasal media (NBM, Gibco 21103-049) supplemented with B27, 10ng/mL BDNF (Peprotech, 450-02), 10ng/mL CNTF (Peprotech, 450-13), and 10ng/mL GDNF (Peprotech, 450-10), ROCK inhibitor, puromycin, and doxycycline. These NGN2-induced neurons were plated on Matrigel coated plates and grown in NBM media containing B27, BDNF, CTNF, GDNF, puromycin and doxycycline. All treatments were carried out on DIV14.

### Method Details

#### Constructs

DsRed-Mito plasmid (Clontech) and shRNA plasmids targeting CLUH (Origene) and SYNJ2BP (pLKO1.1, Sigma), as well as control plasmids were purchased from the respective vendors. The puromycin cassette of pLKO1.1 was replaced with BFP amplified from mito-BFP (Addgene 49151) using BamHI and NsiI restriction enzymes. YFP-Parkin, mito-Raspberry-7, mito-BFP, tagRFP-T-DDX6, AnkyrinG-mCherry, mCherry-Rab7a and mito-Keima were acquired from Addgene (23955, 55931, 49151, 119947, 42566 55127 and 131626 respectively). Plasmids encoding shRNA against SYNJ2 and SYNJ2BP were purchased in pLKO from Sigma (TRCN0000050377 and TRCN0000139049 respectively) as well as a control shRNA plasmid (TR30021). The Puromycin cassette in pLKO was replaced by BFP using restriction enzymes. PINK1 cDNA was purchased from Transomic and inserted into the UBC-pHAGE backbone using NotI and ClaI restriction enzymes, while at the same time adding an EcoRI restriction site downstream of the 3’UTR. In order to insert the 12xMBS-PBS cassette derived from Addgene plasmid 52984, the EcoRI site in the UBC-pHAGE backbone was destroyed by site-directed mutagenesis and the EcoRI digestion product of Pcr4-12xMBS-PBS (Addgene 52984) was inserted into the newly introduced EcoRI site downstream of the PINK1 3’UTR. The correct orientation was verified by sequencing. Beta actin-12xMBS-PBS, Cox4i- 12xMBS-PBS and Atp5f1b-12xMBS-PBS were constructed by replacing the PINK1 sequence with either beta actin, Cox4i or Atp5f1b sequence amplified from rat hippocampal cDNA using NotI and BamHI restriction enzymes. PINK1 kinase dead (KD) mutation was introduced by site-directed mutagenesis. Constructs with portions of PINK1 were derived from the KD mutant by digesting the plasmid with BamHI and replacing the C-terminal part of PINK1 with BFP derived from mitoBFP. Further modifications were achieved using restriction-free cloning (van den Ent and Löwe, 2006). Modification of the split-Venus construct (Addgene plasmid 52985) was performed by Gibson Assembly (Gibson et al., 2009) and included the addition of the rat beta actin zipcode derived from beta-actin-12xMBS-PBS, replacement of the IRES with a P2A ribosomal skip site and removal of the nuclear targeting signal(s). PINK1-N-Kaede and mito-Kaede were generated by insertion of PINK1 Aa 1-624 or Cox8a Aa 1-36 (amplified from mito-Raspberry) in frame before N1-Kaede (Addgene 54726; kind gift from Michael Davidson) using restriction enzyme digestion.

Myc-tagged SYNJ2BP was constructed using mycOmp25-phageNco-forward CTGACccatggacATGGAGCAGAAACTCATCTCTGAAGAGGATCTGAACGGACGGGTGGATTATTTAG and Omp25-phageCla-reverse CTCTAATCGATtcaGAGCTGCTTTCGGTATC primers and inserted into the UBC-pHAGE backbone using NcoI and ClaI restriction sites. A shRNA resistant version was constructed using site-directed mutagenesis to introduce five silent mutations in the shRNA targeting region. FLAG-SYNJ2a and SYNJ1-FLAG were kind gifts from Pietro De Camilli (Nemoto et al., 1997). An outer membrane targeted version of SYNJ2a was constructed using restriction-free cloning to replace the cytosolic part of myc-SYNJ2BP with SYNJ2a, resulting in SYNJ2 (amino acids 1-1218) fused to the SYNJ2BP transmembrane domain (amino acids 110-145). Point mutations in the RRM domain were introduced using site-directed mutagenesis. For expression in HEK cells and for CLIP a shorter version was generated starting at amino acid 880 through restriction-free cloning. For AAV- production, the PINK1-12xMBS-PBS sequence was also inserted in pAAV-MCS (Stratagene) using the XhoI and HindI sites. The control plasmid AAV-GFP has been described before (Park et al, 2008). AAV-particles were produced at Boston Children’s Hospital viral core.

#### Neuronal cultures in microfluidic devices

RD450 microfluidic neuron devices (XONA Microfluidics) were used as described before (Ashrafi et al., 2014). Briefly, the devices were sterilized by spraying with 70% Ethanol and dried in a tissue culture hood. The dry devices were attached to coverslips or 6-well glass bottom plates coated overnight with 20 μg/mL poly-L-Lysine and 3.5 μg/mL laminin that had been washed twice with distilled water and left to dry for 2-3 min under the hood. Dissociated hippocampal neurons were pelleted at 1500 g for 4 min, resuspended at a final concentration of 20×10^3^/ µl in NB+PSG+B27. 5 µl were plated into one of the somal compartments and incubated at 37°C in 5%CO2 for 15 min before filling up the wells with NB+PSG+B27. 50% of the medium was replaced every 2-3 days with fresh NB+PSG+B27.

#### Mitophagy detection

##### Parkin translocation

Rat hippocampal neurons in microfluidic devices were transfected for four hours on DIV6 with mito-dsRed and YFP-Parkin using lipofectamine 2000 transfection reagent (Thermo Fisher) in medium lacking B27. On DIV8 cells were incubated in Hibernate E (BrainBits) with or without 70µM Cycloheximide (Sigma) for 4h, before live cell imaging at a spinning disk microscope (Yokogawa CSU-X1, Olympus IX81) equipped with an electron-multiplying charge-coupled device camera (Andor iXon; Oxford instruments) using a 40×/NA 1.3 oil immersion lens and Metamorph software (Molecular Devices). Images were taken before and 20 min after addition of 40 µM Antimycin A (Sigma) in the axonal chamber, leaving all settings identical, including detector sensitivity and camera exposure time.

##### Mito-mKeima mitophagy index

Mouse hippocampal neurons were seeded in 24-well glass bottom plates (CellVis) at a density of 100*10^3^ and maintained as described above. On DIV6 Neurons were transfected for 20min with mito-mKeima and shRNA against SYNJ2BP or Control shRNA using lipofectamine 2000 transfection reagent (Thermo Fisher) in medium lacking B27. On DIV9 cells were incubated in Hibernate E (BrainBits) with or without 40µM Antimycin A (Sigma) for 1h before live-imaging at the Imaging Facility of Max Planck Institute of Biochemistry, Martinsried, on a LEICA (Wetzlar, Germany) SP8 FALCON confocal laser scanning microscope equipped with a HCX PL APO 63x/1.2 motCORR CS water immersion objective. Keima green was excited at 442 nm and Keima red at 550 nm. Emission was detected sequentially from 555-620 nm for both excitation wavelengths. Imaging settings were kept constant for all conditions.

#### RNA live cell imaging

Rat or mouse hippocampal neurons were seeded in 24-well glass bottom plates (CellVis) at a density of 100*10^3^ and maintained as described above. On DIV5-7, cells were washed three times with NB+PSG and transfected using lipofectamine 2000 transfection reagent (Thermo Fisher) for 20-40 min. After transfection, the original conditioned medium with B27 was returned after three washes. The ideal ratio between the construct encoding the split-Venus parts and the construct encoding the mRNA with the respective binding sites was determined empirically to be approximately 1:4. While we typically observed a high amount of co-transfection in our cultures (around 96%, compare Wang et al., 2011), only around 10% of the cells that were transfected with the mitochondrial marker also showed successful fluorophore reconstitution. In addition, we had to transfect with 2-4µg DNA per well in order to observe export of the construct from the nucleus. Constructs were expressed for 1-2 days, except in the case of cells cotransfected with shRNA, which were imaged after 3 days to provide enough time for effective reduction of the protein. Imaging was performed in Hibernate E with a spinning disk microscope; using either a Yokogawa CSU-X1 (Olympus IX81) equipped with an electron-multiplying charge-coupled device camera (Andor iXon; Oxford instruments) using a 40×/NA 1.3 or 60×/NA 1.2 oil immersion lens and Metamorph software (Molecular Devices), or a Eclipse Ti2 spinning disk microscope (Nikon) equipped with a DS-Qi2 high-sensitivity monbochrome camera (Nikon) using a 60×/NA 1.2 oil immersion lens and NIS-elements software (Nikon). For puromycin treatment, puromycin was added at a concentration of 200 µg/mL to the medium 1 h prior to imaging.

COS-7 cells were maintained in DMEM + GlutaMax supplemented with penicillin/streptomycin (Life Technologies), and 10% FBS (Atlanta Premium) and transfected and imaged as described for neurons. Expression of the IRES construct was not as efficient as the P2A split Venus construct and we therefore used the P2A construct for the experiments in COS-7 cells.

#### *In situ* hybridization and immunocytochemistry

RNAscope hybridization was performed as described in Cosker et al., 2016. Briefly, mouse hippocampal neurons were grown on glass coverslips and fixed on DIV7 with 4% paraformaldehyde for 15 min. After dehydration in a dilution series of ethanol, cells were stored at -20°C for up to one month. Cells were rehydrated and permeabilized with 0.1% Tween 20/PBS for 10 min, rinsed in PBS and incubated in a 1:5 dilution of Protease III (ACD) for 10-20 min at 40°C in a preheated hybridization oven. *In situ* hybridization with mouse PNK1 probes (ACD) was performed at 40 °C for 2h and the detection reactions were performed according to the manufacturer’s instructions.

For immunodetection, coverslips were rinsed with PBS and blocked with PBS/0.3% Triton X/4% goat serum for 1h at RT. Samples were incubated with primary antibodies for 2-3 hours at RT, washed twice fast and twice for 10 min at RT with PBS/0.3% Triton X before incubation with Alexa488 or Alexa647-conjugated secondary antibodies in PBS/0.3% Triton X for 2h at RT. After washing in PBS/0.3% Triton X coverslips were mounted in Fluoromount G (Southern Biotech) and imaged at a confocal microscope (LSM710, Carl Zeiss) using a 63x/NA 1.4 oil immersion objective and ZEN 2009 software (Carl Zeiss) or at a widefield microscope (EVOS M5000, Thermo Fisher) using a 10x objective.

STED imaging was perfomed on RNAscope samples using the Opal570 (RNAscope) and Abberior635P (Immunostaining) fluororescent probes using the same protocol and imaged at a Stedycon system (Abberior) mounted an a Leica DMRXA2 body, using a 100x/NA 1.4 Oil immersion objective and a 775 nm STED laser.

#### Kaede photoconversion

Hippocampal neurons were seeded in 24-well glass bottom plates at a density of 100*10^3^. On DIV7, cells were transfected with PINK1-N-Kaede or mito-Kaede using lipofectamine 2000 transfection reagent for 20 min. Constructs were expressed for 48 hrs to provide enough time for effective expression of the protein. Imaging was performed in Hibernate E with a WF2 Leica Thunder microscope using a HC PL APO 63x/1.20 WATER UV objective and LAS X software. Prior to photoconversion, a defined region of the axon containing Kaede-green fluorescent mitochondria was imaged using 488 and 558 nm multicolor-illumination. Immediately thereafter, the Kaede-green mitochondria were photoconverted to Kaede-red by using a 405 nm laser scanner while visually assessing for residual green fluorescence. The axonal regions were re-imaged using 488 and 558 nm multicolor-illumination directly after and 45-60 min post-photoconversion. Only Kaede-red mitochondria, which were still within the defined region, were used for analysis.

#### Lentiviral transduction

Lentiviral particles were produced in HEK293T cells as described previously (Pekkurnaz et al., 2014). Hippocampal neurons were transduced on DIV1 or 2 and lysed for Western Blot analysis after 4 days. Infection rates were 60-90%. Western blotting was performed using standard procedures and blots were decorated with the following antibodies diluted in PBS + 5% milk: Mouse anti-beta-actin Monoclonal Antibody (AC-74) (1:1000, Sigma), Mouse monoclonal anti-Glyceraldehyde-3-PDH (GAPDH) antibody (1:1000, EMD Millipore), Rabbit Polyclonal Anti-CLUH Antibody (1:250, Origene), Rabbit polyclonal anti-SYNJ2BP antibody (1:200, Proteintech). Western Blot analysis was performed using LI-COR secondary antibodies and an Odyssey CLx Infrared Imaging System (LI-COR Biosciences).

#### RNA isolation and qRT-PCR

For analysis of RNA abundance after CLUH knockdown, cells were harvested 4 days after lentiviral transduction and RNA was isolated using the QIAGEN RNeasy Mini Kit. cDNA was generated using the qScript™ cDNA SuperMix (Quantabio) and a qPCR assay was performed using PerfeCTa SYBR® Green FastMix (Quantabio) in a StepOnePlus Real PCR machine (Thermo Fisher). Abundance was calculated relative to beta actin and control shRNA using comparative Ct using the formula: 2^-ΔΔCt^ (relative quantitation) from 3 independent biological repeats.

For analysis of axonal transcripts, rat hippocampal neurons were grown in microfluidic devices. On DIV7 the axonal and soma chamber were lysed individually and RNA was isolated using the QIAGEN RNeasy Mini Kit. cDNA was generated using the qScript™ cDNA SuperMix (Quantabio) and a qPCR assay was performed using PerfeCTa SYBR® Green FastMix (Quantabio) in a StepOnePlus Real PCR machine (Thermo Fisher). Abundance was calculated relative to mitochondrial S12 rRNA and the soma chamber using comparative Ct using the formula: 2^-ΔΔCt^ (relative quantitation) from 3 independent biological repeats.

For comparison of expression of SYNJ2a and RRBP1 in hippocampal neurons and fibroblasts, DIV7-9 neurons and low passage primary fibroblasts were harvested and RNA was isolated using the QIAGEN RNeasy Mini Kit. cDNA was generated using the qScript™ cDNA SuperMix (Quantabio) and a qPCR assay was performed using PerfeCTa SYBR® Green FastMix (Quantabio) in a StepOnePlus Real PCR machine (Thermo Fisher). Abundance was calculated relative to beta actin using standard curves generated from beta actin, SYNJ2a and RRBP1 constructs (absolute quantitation), using three to five independent samples.

#### Retinal AAV-injection and RNA isolation

Surgical procedures were performed as described in Park et al. (2008). Mice were anaesthetized with ketamine and xylazine and, to protect the cornea during surgery, eye ointment containing atropine sulfate was applied. A glass micropipette was inserted at an angle posterior to the ora serrata to avoid damage to the lens. 1 µl AAV2-GFP and AAV2-PINK1- 12xMBS-PBS of similar titers were injected intravitreally. As post-operative analgesic mice received Buprenorphine (0.1 mg/kg, Bedford lab) for 24 hr. After four weeks mice were sacrificed and both the retina and the optic nerve prior to the optic chiasm were collected and stored in RNAlater (QIAGEN) at 4 °C overnight. After removal of RNAlater reagent, 500µl TRIZOL was added and the tissue homogenized on ice using a micropestle mixer. After 5min incubation at RT, 100 µl Chloroform was added and the mixture was vortexed for 15 sec and incubated for 2 min at RT. Phase separation was achieved during centrifugation at 12000 g for 15 min at 4°C. The aqueous phase was collected and the RNA precipitated with 500 µl Isopropanol for 15 min at RT. RNA was pelleted during a spin at 12000 g for 15min, washed with 70% ethanol, and resuspended in 50 µl RNase-free water for retina - 25µl for optic nerve samples. cDNA was generated using the qScript™ cDNA SuperMix (Quantabio) and a qPCR assay was performed using PerfeCTa SYBR® Green FastMix (Quantabio) in a StepOnePlus Real PCR machine (Thermo Fisher). Abundance was calculated relative to beta actin and the retinal amount using comparative Ct using the formula: 2^-ΔΔCt^ (relative quantitation). Four retina/optic nerve pairs were analyzed per transcript.

#### Cross-Linking Immunoprecipitation (CLIP)

HEK293T cells were grown in 6 well plates and transfected with 3 µg/well Myc-SYNJ2aRRM- mito or its VQL/AAA mutant. UV irradiation was performed by washing the cells with PBS and placing the plate on ice in a CL-1000 crosslinker (UVP) and exposing the plate to 400 mJ/cm^2^ 254 nm UV light. After irradiation, cells were harvested in Lysis Buffer (1% Triton, 20mM Tris pH 7.4, 200mM NaCl, RNAsin (1:100 Promega), Protease inhibitor cocktail III (Millipore) and 200 µM PMSF) and cleared by centrifugation at 12000 g for 1 min. The supernatant was incubated with 3 µl anti-myc antibody (mouse 9E10, Novus) /mL lysate for 1h at 4°C. ProteinA sepharose beads were blocked with 3% BSA in lysis buffer for 30min, washed with PBS and added to the lysate. After 30 min incubation at 4°C beads were collected by centrifugation at 2000g for 30sec and washed three times with lysis buffer. Samples were eluted by addition of Laemmli Buffer and boiling at 95°C for 3 min prior to analysis by gel electrophoresis and immunoblotting with Rabbit myc-tag antibody, 71D10 (Cell Signaling).

#### Proximity ligation assay (PLA)

The proximity ligation assay was performed according to the manufacturer’s instructions (Sigma-Aldrich). Briefly, primary mouse hippocampal neurons were grown on glass coverslips, fixed on DIV 8 with 4 % paraformaldehyde for 15 min and permeabilized with 0.3 % Triton/PBS for 10 min followed by a 1 h incubation with Duolink blocking solution at 37 °C. Neurons were incubated with primary antibodies (SYNJ2BP-SYNJ2 interaction, SYNJ2BP-ATP5B interaction, SYNJ2-RHOT1 interaction; mouse polyclonal SYNJ2BP antibody, 1:50, Sigma-Aldrich; rabbit polyclonal SYNJ2 antibody, 1:50, Proteintech; rabbit polyclonal ATP5b antibody, 1:600, Sigma-Aldrich; mouse monoclonal RHOT1 antibody (4H4), 1:500, Sigma-Aldrich) diluted in Duolink antibody diluent at 4 °C overnight. Neurons were washed two times with Buffer A (0.01 M Tris, 0.15 M NaCl and 0.05 % Tween 20) at RT for 5 min, incubated with Duolink PLA Probes (Anti-Rabbit Plus and Anti-Mouse Minus) at 37 °C for 1 h, again washed two times with Buffer A at RT for 5 min, then incubated with Duolink ligation solution at 37 °C for 30 min, again washed two times with Buffer A at RT for 5 min and incubated with Duolink amplification solution at 37 °C for 100 min. After two washes with Buffer B (0.2 M Tris, 0.1 M NaCl) at RT for 10 min and a final wash with 0.01x Buffer B for 1 min the coverslips were mounted in Fluoromount G (Invitrogen) and imaged at a Nikon Ti2 spinning disk microscope using a 60x/NA 1.40 oil immersion objective. For puromycin treatment, puromycin was added at a concentration of 200 µg/ml to the medium 1 h prior to fixation. The number of PLA puncta per soma was quantified and normalized to the number of the SYNJ2BP-SYNJ2 PLA puncta.

### QUANTIFICATION AND STATISTICAL ANALYSIS, SUPPLEMENTAL INFORMATION

Throughout the paper, data are expressed as mean ± SEM. Statistical analysis was performed with Excel (Windows) or R (The R foundation) using student’s t-test for gaussian distributions. When comparing multiple conditions a one-way ANOVA test for statistical significance was followed up by a Bonferroni post-hoc test. p < 0.05 was considered significant (*), with further levels defined as p<0.01 (**), p<0.001 (***) and p<0.0001 (****). Where practical, especially for values >0.01, actual p values are given in the figure or figure legend.

Quantification of Western Blots was performed in Image Studio Lite (LI-COR) using the local background correction. Quantification of microscopy data was performed using Image J. Co-localization was analyzed in z-stack images using the JaCOP and “straighten” plugins as described in Graber et al. (2013). For dendrites and axons imaged with the split-Venus approach, after maximum z projection neurites were straightened with a 20 px margin. For the cell body quantification, a 10 by 10 µm square was chosen within the cell body and no z projection was performed. The position of the square was chosen based on the mitochondrial signal to exclude the nucleus and blinded to the phenotype of the RNA channel. Mander’s correlation coefficients were exported to Excel and plotted in R using the R boxplot function. Box and whisker plots represent the median (line), 25th-75th percentile (box) and 10th-90th percentile (whiskers).

Time-lapse imaging was performed by imaging every 1 sec for 90 sec. Movies were analyzed using Kymolyzer macro for ImageJ developed in the laboratory (Pekkurnaz et al., 2014). Time spent in motion was averaged for every dendrite separately, which creates a gaussian distribution of average time spent in motion per neurite.

For the histogram of length traveled, 43 movies that showed at least one moving PINK1 RNA particle were converted to kymographs and the distance in x was measured for the intervals of movement. The corresponding mitochondrial kymograph was then examined and, if a similar mitochondrial track was observed, the even was scored as a co-movement.

Keima images were analyzed using Image J. Mitochondrial were identified on a thresholded image using the particle analyzer and the intensities of the non-thresholded images were calculated. For each field of view the mean integrated density was exported to Excel. The mitophagy index was expressed as in^9^ by calculating the ratio of the integrated density signals [Keima red/(Keima red + Keima green)].

Kaede images were analyzed using Image J. Mitochondrial were identified on a thresholded image using the particle analyzer and the intensities of the non-thresholded images were calculated. The mean intensities as well as a background intensity were exported to Excel and the substracted values were visualized with the R ggplot2 library.

Myc-SYNJ2 ICC images were analyzed using ImageJ. Myc signal was identified on a thresholded image using the particle analyzer and the intensities of the non-thresholded images were calculated. The mean integrated densities of the myc signal and the mito-mRaspberry signal were exported to Excel. For each neuron, the mean integrated density of the myc signal was normalized to the mean integrated density of the mito-mRaspberry signal. HA-splitVenus ICC images were analyzed using ImageJ. Axons were traced using the segmented line tool for each field of view and the mean integrated densities of the HA and iRFP signals were measured and exported to Excel. The HA intensity was normalized to the iRFP intensity. The first two fields of view are classified as “proximal”, whereas all subsequent fields of view were categorized as distal as they are more than 750 µm away from the cell body.

